# Tunable Universal OR-gated CAR T cells for AML

**DOI:** 10.1101/2024.04.13.589307

**Authors:** Menna Y. Siddiqui, Jingyao Chen, Madeline Loffredo, Seunghee Lee, Han Deng, Yongshuai Li, Nelia Leemans, Tim Lu, Brian S. Garrison, Marcela Guzmán Ayala, Nicholas W. Frankel, Wilson W. Wong

**Affiliations:** Department of Biomedical Engineering and Biological Design Center, Boston University, Boston, MA; Department of Materials Science and Engineering, Boston University, Boston MA; Senti Biosciences, South San Francisco, CA

## Abstract

Acute myeloid leukemia (AML) is a hematopoietic malignancy characterized by antigen heterogeneity and poor prognosis. A potential therapeutic approach to address this heterogeneity is targeting multiple surface antigens to prevent antigen escape and relapse. Chimeric antigen receptor (CAR) T cells are an adoptive cell therapy that have demonstrated remarkable clinical success in the treatment of B cell malignancies, and many efforts are underway to adapt them to myeloid malignancies. To tackle the heterogeneity of AML, logically targeting multiple antigens through an “A OR B” gated CAR circuit would be desirable. Here we combined FLT3 antigen targeting with the well characterized CD33 myeloid marker as a combinatorial OR gate approach using our split, universal, programmable (SUPRA) CAR platform. The split platform affords tunability over activation levels and multiplexed targeting that cannot be achieved through a tandem bispecific approach. We systematically characterized the specificity and sensitivity of different SUPRA CAR adapters against each target individually and in combination against a panel of target cell lines. Our results demonstrate that this CAR system can effectively target two antigens with equivalent efficacy to conventional CARs while reducing the engineering burden associated with designing CAR T cells against multiple antigens. Furthermore, we can characterize an effective dose range where off-target cytotoxicity against hematopoietic stem and progenitor cells is minimized. With the recent clinical advances in universal CAR designs, our SUPRA OR gate has the potential to provide an effective and safer solution to treating AML.

## Introduction

Acute myeloid leukemia (AML) is a subset of leukemia that begins in the blood-forming cells of the bone marrow, and has a particularly poor prognosis, with a 5-year survival rate of just over 30%^1–3^. Despite advances in conventional treatments such as chemotherapy and stem cell transplantation^4–8^, there is a significant unmet need for new and effective therapies for AML patients^9,10^. Immunotherapy using T cells engineered to target cancer-specific surface antigens has emerged as a promising approach for cancer treatment. Chimeric Antigen Receptor (CAR) T cell therapy has demonstrated remarkable efficacy against certain hematological malignancies, with six FDA approved clinical products targeting CD19 in B-cell malignancies and BCMA in multiple myeloma^11–16^. Additionally, there are over 500 clinical trials currently underway investigating CAR T cell treatment for other diseases, including AML^17,18^.

CAR T therapy involves genetically modifying a patient’s own T cells to express a CAR that recognizes and binds to a specific antigen present on cancer cells. Once the CAR T cells are infused back into the patient, they can selectively target and kill cancer cells, leading to durable and potentially curative responses. However, CAR T therapy for AML has been challenging due to the heterogeneity of this disease – it is characterized by a wide spectrum of mutations that influence the phenotype and disease progression for each individual patient (**Fig 1A**)^19–21^. In addition to clonal heterogeneity, many of the surface antigens shown to be overexpressed in AML are also expressed on healthy hematopoietic stem cells (HSCs), which has made it difficult to find a suitable antigen target for therapy (**Fig 1B**)^22^.

**Figure 1.**
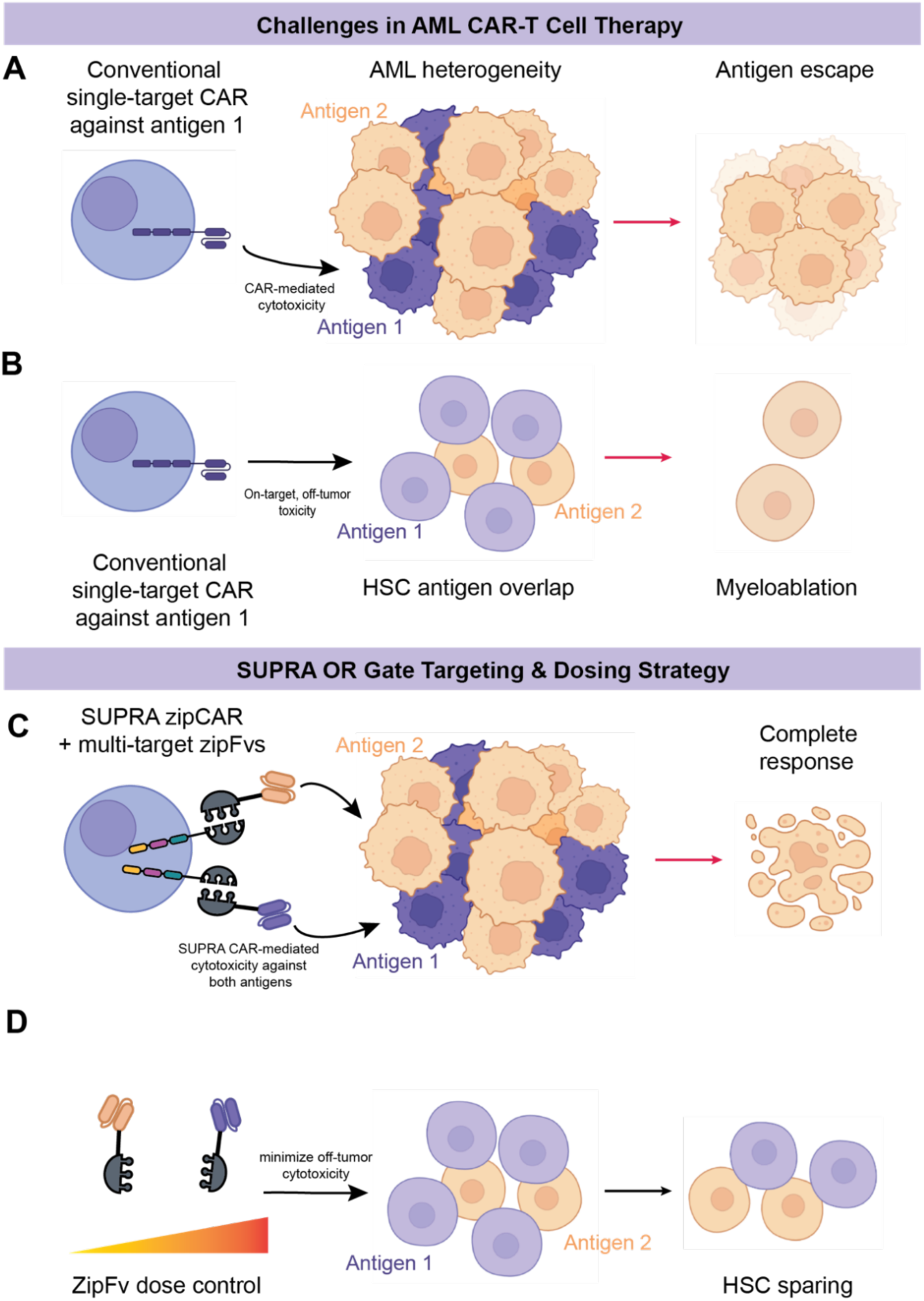
SUPRA can address common challenges in AML CAR-T cell therapy. A. AML heterogeneity poses a challenge for conventional CAR T therapy against a single target, leading to antigen escape. B. Healthy hematopoietic stem cells express many of the same antigens as AML, leading to myeloablation. C. By targeting multiple antigens using a SUPRA OR-gate configuration, we can combat antigen escape and approach a complete response. D. By optimizing the dosing of zipFv, we can identify a therapeutic dose that spares HSCs while eliminating AML cells.

Current CAR T therapies, including many of those in clinical trials, use conventional CAR designs consisting of a fixed, antigen-specific scFv targeting a single cancer antigen coupled with intracellular signaling domains for T cell activation. For B-cell malignancies, CD19 has proven to be an ideal target antigen due to its restricted expression on B-cell lineages and not on other hematopoietic lineages. Since no such single antigen has been identified for AML, conventional CAR designs have shown severe on-target, off-tumor toxicity^23^. Moreover, the heterogeneity of AML makes it difficult for a single-target strategy to achieve a complete response without the risk of refractory disease.

Combinatorial antigen-targeting presents a potentially effective strategy for addressing heterogeneity and off-tumor toxicity. Our group and others have implemented engineered Boolean logic gates, such as “OR”, “AND”, and “NOT”, to advance the targeting capabilities of CAR T cells^24–33^. In particular, OR gate logic can target multiple antigens to prevent antigen escape and subsequent disease relapse due to AML heterogeneity (**Fig 1C**). Furthermore, by decoupling the antigen-targeting modality from the CAR design, a split CAR design could add an additional layer of safety and tunability not available within other approaches^34^.

To achieve this, we have previously developed a split, universal, and programmable (SUPRA) CAR system composed of a universal chimeric receptor expressed on the T cell, zipCAR, coupled with a soluble antigen-binding adapter, zipFv, that contains a leucine zipper and scFv^35^. The SUPRA system enables OR gate logic and a tunable T cell response that corresponds to the concentration of each zipFv supplied to the system. We leverage this design to target two AML-associated antigens, CD33 and FLT3, which have been identified as therapeutic targets and are currently being explored in single-target conventional CAR T clinical trials^23,36,37^. Over 80% of AML cases are CD33+, and the first targeted AML therapy introduced was gemtuzumab ozogamicin, a monoclonal anti-CD33 antibody conjugated with an anti-tumor drug^38,39^. However, fatal adverse events such as infection and acute respiratory distress syndrome led to its temporary withdrawal for 7 years before re-approval^40^, and it remains efficacious mainly in patients with very high expression of CD33. Recently, Bispecific T-cell Engagers (BiTEs) and anti-CD33 CAR T designs have been explored in Phase I clinical trials^41,42^, with mixed results due to rapid disease progression and the manufacturing challenges of these modalities. Similarly, the FMS-like tyrosine kinase-3 (FLT3) is one of the most frequently mutated genes in AML, accounting for approximately 30% of AML cases^3,22,43,44^. This mutation is additionally correlated with poor outcomes, and there remains to be an FDA-approved targeted therapy against this antigen. CAR-T designs against FLT3 are being actively explored^36,45^, making it a suitable candidate for our proposed combinatorial approach. By decoupling the targeting modality from the CAR T design, we anticipate that our SUPRA system can alleviate some of the manufacturing challenges of validating new CAR-T therapies while allowing for control over dosing schedules to obtain optimal efficacy for each patient. We show that the relative affinity of each scFv to its target antigen has significant effects on the efficacy of combinatorial dosing, which we have characterized to identify an optimal dose for each target antigen. Additionally, we demonstrate how zipFv dose control can identify a therapeutic window that effectively eliminates tumor cells with varying antigen expression levels while sparing most healthy HSCs (**Fig 1D**).

## Results

### Design and Characterization of SUPRA CAR against target antigens FLT3 and CD33

The SUPRA system comprises a universal zipper CAR (zipCAR) and a recombinant protein combining a leucine zipper and an scFv (zipFv). To design zipFvs targeting FLT3 and CD33, we chose two clonal variations of each targeting antibody fragments used in the scFv design: anti-FLT3 (NC7), anti-FLT3 (D4-3), anti-CD33 humanized monoclonal 195 (hu-195), and anti-CD33 Mylotarg (mylo) (**Fig 2A**). These were fused with an EE zipper through a GS linker and paired with a third-generation RR-CAR (**Supplemental Fig 1A**). To characterize the function of these zipFvs, we screened all zipFvs with RR-CAR transduced NFAT-GFP Jurkat T cells (**Fig 2B**) against engineered CD33+, FLT3+, and FLT3+CD33+ K562 target cell lines in *in-vitro* co-culture activation assays. Briefly, RR-CAR transduced NFAT GFP cells were co-cultured with target cells at a 1:1 effector to target (E:T) ratio with or without 50 ng of adaptor zipFv, and NFAT transcriptional activity was measured via the T cell activation NFAT-GFP transcription reporter expression to assess on-target and off-target activity. The FT3 NC7 zipFv variant showed significantly higher activation than the D4-3 variant, while both showed higher activation levels than the off-target CD33 zipFvs when targeting FLT3+ K562 (**Fig 2C**). In contrast, both CD33 zipFv variants had high on-target activity against CD33+ K562 that was significantly higher than off-target activity by FLT3 zipFv variants (**Fig 2D**). Similar to the results against single positive K562 targets, FLT3 NC7 variant showed significantly higher activity compared to D4-3 against CD33+FLT3+ K562 (**Fig 2E**). Although the CD33 mylo variant showed slightly higher activity than the hu195 variant against the double+ target cells, the protein production yield of CD33 hu195 zipFv was significantly higher (**Supplemental Figure 1B**), leading us to favor this variant over the CD33 mylo zipFv.

**Figure 2.**
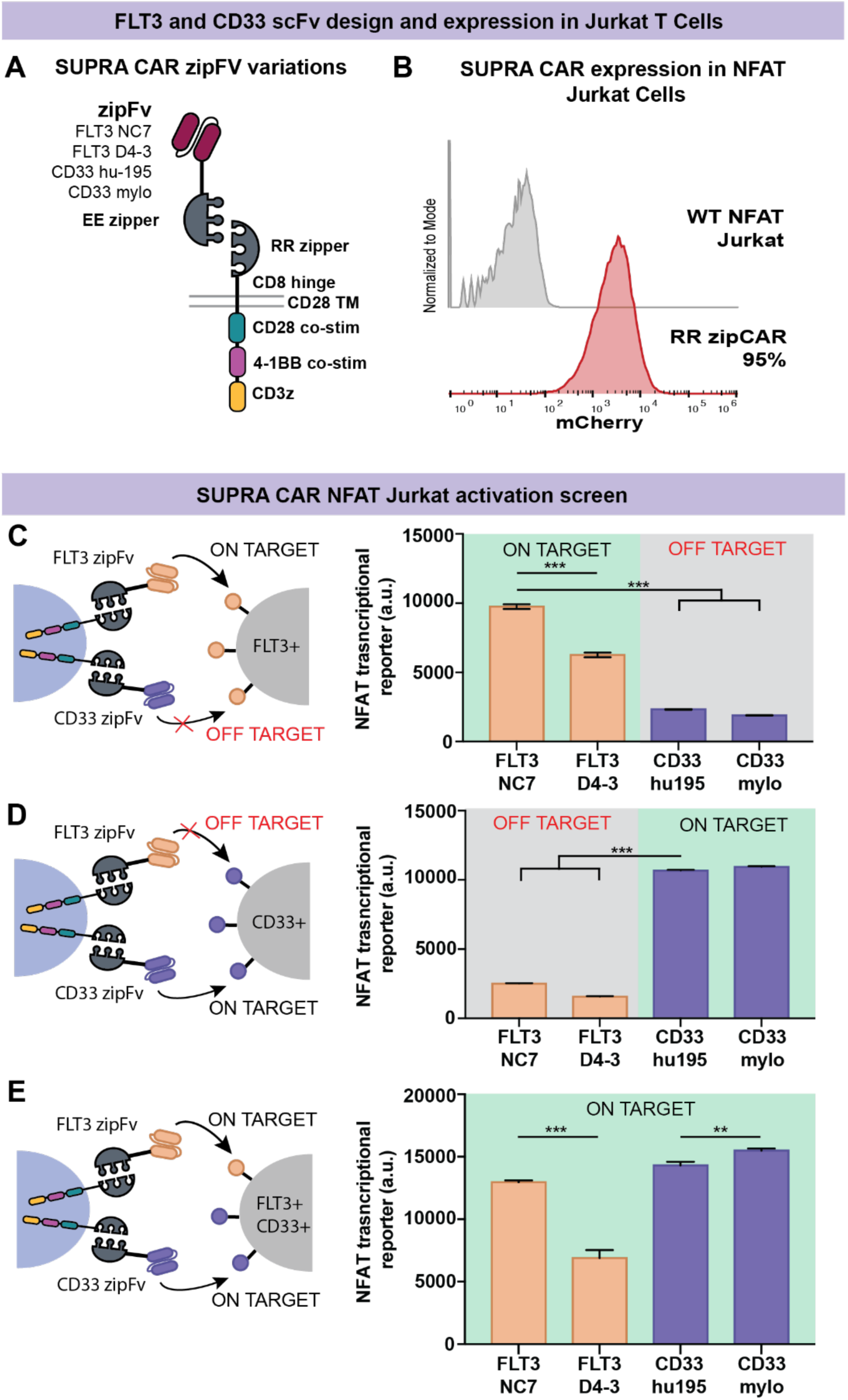
SUPRA CAR OR-Gate against FLT3 and CD33 design and initial activity screening. A. Four zipFv variations were designed as a fusion of an EE leucine zipper to FLT3 or CD33 scFv clones and paired with a 3^rd^ generation RR CAR containing both CD28 and 4-1BB costimulatory domains in addition to the CD3z activation domain. B. RR CAR was transduced into NFAT-GFP Jurkat T cells with a transduction efficiency of 95%. C. FLT3+ K562 cells were co-cultured with RR CAR in the presence of either FLT3 zipFv to assess on-target activity or CD33 zipFv to assess off-target activity. D. CD33+ K562 cells were co-cultured with RR CAR in the presence of either CD33 zipFv to assess on-target activity or FLT3 zipFv to assess off-target activity. E. Double+ CD33+/FLT3+ K562 were co-cultured with RR CAR in the presence of each zipFv variation to assess on-target activity. In all co-culture conditions, an E:T ratio of 1:1 was used, and 50 ng of the corresponding zipFv was added. CAR-T activation is reported as the expression of GFP transcribed downstream of the NFAT reporter. Values shown are the means of technical triplicate samples with error bars indicating +1 standard deviation, and p-values were calculated as described in the methods.

In addition to the EE-RR zipper SUPRA design, we also characterized the activity of the SUPRA system generated with FOS leucine zippers, pairing with a JunD zipCAR transduced into NFAT-GFP Jurkat cells (**Supplemental Fig 2A**). However, both FLT3 FOS zipFvs show reduced on-target activity that was not significantly higher than the off-target signal (**Supplemental Fig 2B**). In contrast, the CD33 FOS zipFv variants showed similarly high on-target activity compared to their EE counterparts (**Supplemental Fig 2C**). We additionally designed conventional single-target CAR constructs for each scFv variant **(Supplemental Fig 1A**). We transduced these constructs into NFAT-GFP Jurkat T cells to compare their activity against our SUPRA system (**Supplemental Fig 3A**). While the activation trends were similar, all conventional designs exhibited lower maximum activation levels against their corresponding single+ targets, resulting in a lower fold change between on-target and off-target states (**Supplemental Fig 3B-C**). Based on results of these NFAT Jurkat activation assays, the clonal variation FLT3 NC7 outperformed the FLT3 D4-3 clone, and the EE-RR leucine zipper system outperformed the FOS-JUN leucine zipper pairs for FLT3 targeting. Importantly, protein production yield varied significantly between clonal variations, with FLT3 NC7 and CD33 hu-195 yielding more zipFv protein. Thus, we decided to move forward with these two zipFvs and the EE-RR leucine zipper pairs for further characterization of the OR gate.

### Primary CD3+ CAR T cell cytotoxicity against AML target cell lines

To assess specific lysis of CD33 and FLT3 expressing target cells, RR-CAR was transduced into primary CD3+ T cells with a transduction efficiency of over 80% for use in *in-vitro* co-culture cytotoxicity assays (**Fig 3A**). As a model for antigen expression heterogeneity, we characterized the antigen expression levels of the engineered K562 cell lines, as well as the MV4-11 human AML cell line and THP-1 acute monocytic leukemia cell line, using a standardized fluorochrome-conjugated bead kit to determine antibody binding capacity against FLT3 and CD33 (**Fig 3B**). Given the variation in expression levels, we determined these cell lines were an acceptable model to test the functionality of the OR gate, with the WT K562 cell line serving as a low-expression negative control.

**Figure 3.**
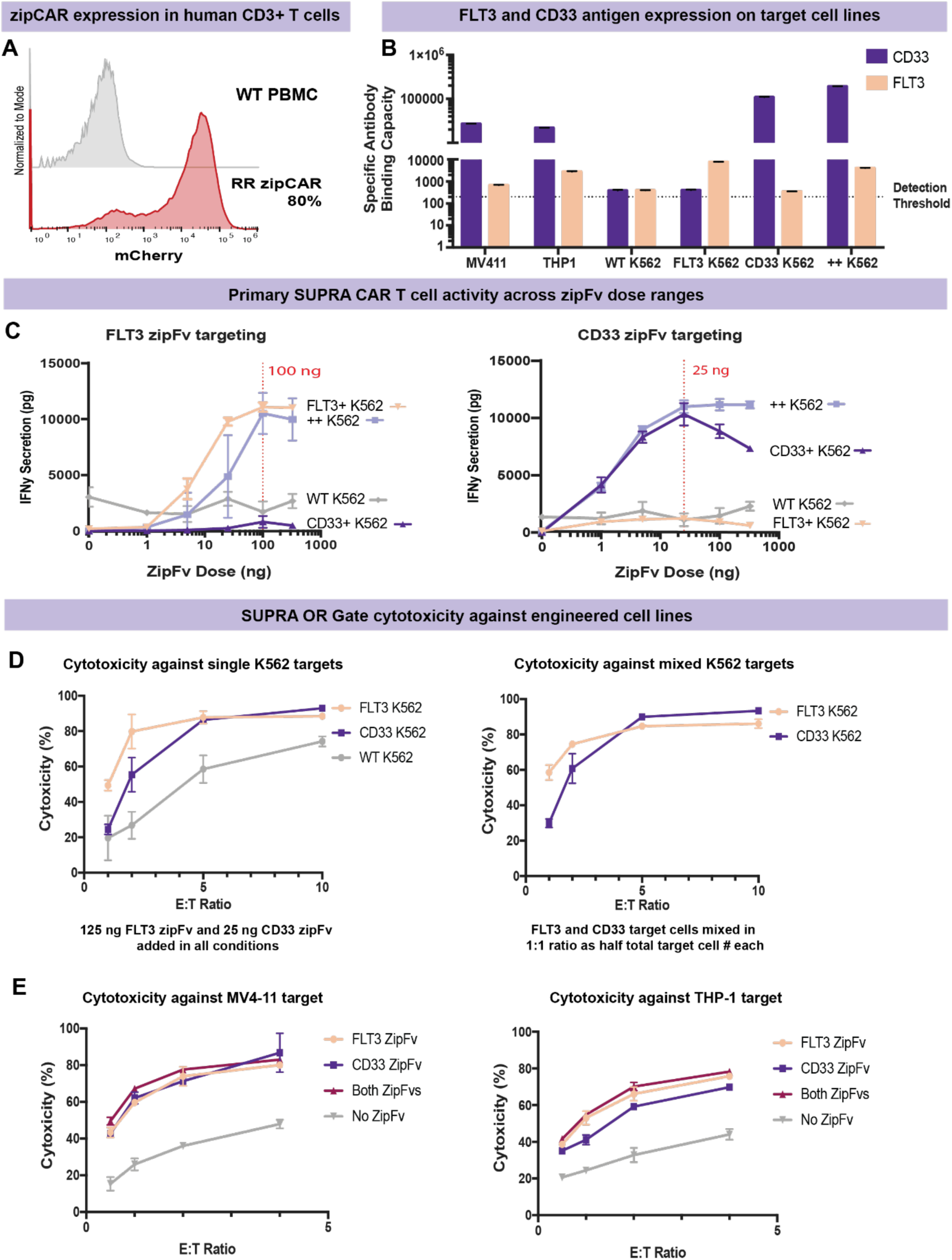
Primary CD3+ SUPRA CAR T cell activity and cytotoxicity against AML target cell lines. A. RR zipCAR was transduced into primary CD3+ human T cells with a transduction efficiency of ∼80%. B. MESF quantitative fluorescence kit from Bangs Laboratories was used to quantify the antibody binding capacity against FLT3 and CD33 antigens on engineered K562 cell lines and MV4-11 and THP-1 cell lines. The QuickCal pre-formatted template was used to plot the standard curve and determine the detection threshold. C. To determine the optimal zipFv dose for each target, RR CAR and target cells were co-cultured at an E:T of 2:1 with varied doses of zipFv, followed by an IFNy ELISA of the supernatant collected 24 hours later. A dose of >100 ng of FLT3 zipFv and 25 ng of CD33 zipFv resulted in maximal activation. D. Using 125 ng of FLT3 zipFv and 25 ng of CD33 zipFv, cytotoxicity against each single positive K562 target as well as an equal mix of both targets, was determined across a range of E:T ratios to confirm effective killing at these doses. E. Cytotoxicity against AML cell lines MV4-11 and THP-1 was also determined using the same zipFv doses across a range of E:T ratios. Values shown are the means of technical triplicate samples with error bars indicating +1 standard deviation.

A key feature of the SUPRA system is zipFv dose-dependent activity, which allows us to choose an optimal dose for maximum T cell activation. To identify this dose for each zipFv, primary CD3+ RR-CAR T cells were co-cultured with K562 target cell lines in a 2:1 E:T ratio with varying amounts of zipFv for 24 hours, followed by the collection of the supernatant to measure IFNy secretion as an indication of T cell activity. The FLT3 zipFv reached maximum activity at a dose of 100 ng in a 200 uL coculture, with minimal off-target activity against CD33+ and WT K562 cells (**Fig 3C**). In contrast, the CD33 zipFv saturated activity at a lower dose of 25 ng and exhibited the Hook effect at higher doses against the CD33+ K562 target (**Fig 3C**). This activity can be explained by differences in the binding affinity of the scFvs against their target antigen, which we further characterize below.

The cytotoxicity against the target cell lines at different E:T ratios was characterized through *in-vitro* co-culture cytotoxic assays using an FLT3 zipFv dose of 125 ng and a CD33 zipFv dose of 25 ng in all experiments to ensure that the presence of both zipFvs did not interfere with the lysis of single positive targets. To calculate specific lysis, target cells were pre-stained with CellTrace Violet to differentiate them from CAR T cells in the downstream flow cytometry analysis (**Supplemental Fig 4**). Both FLT3 and CD33 zipFvs can induce high SUPRA CAR T cell cytotoxicity when combined with single FLT3+ K562 and CD33+ K562 in an E:T dependent manner, respectively (**Figure 3D**). Importantly, at our optimized dosages, we observed significantly higher IFNy secretion from the SUPRA system in on-target conditions when compared to conventional CARs co-cultured at the same E:T ratio (**Supplemental Fig 5A-B**). This indicates that our dose optimization can also maximize CAR activation compared to constitutively active CAR designs. In addition, to confirm that the FLT3 and CD33 SUPRA CAR system can function as an OR gate against two single positive populations of target cells simultaneously, we mixed FLT3+ K562 and CD33+ K562 targets in a 1:1 ratio as half the total target cell number with the addition of the corresponding targeting zipFv, and observed that both zipFvs could successfully trigger SUPRA CAR T cells cytotoxicity against their corresponding target in the mixed target coculture (**Fig 3D**).

We further tested this system against the cell line MV4-11, an AML cell line derived from the blast cells of a 10-year-old male with biphenotypic B-myelomonocytic leukemia that carries the FLT3-ITD mutation, and THP-1, a human monocytic leukemia cell line. Both cell lines have high CD33 expression and detectable amounts of FLT3 antigen expression, and thus are expected to induce cytotoxicity for FLT3 and CD33 SUPRA systems (**Figure 3B**). MV4-11 or THP-1 cells were mixed with FLT3, or CD33 zipFv alone or together, and co-cultured with SUPRA CAR T cells for 24 hours under different E:T ratios. As expected, SUPRA CAR T cells showed high target cell killing efficiency when either or both zipFvs were present that increased in an E:T dependent manner (**Fig 3E**), as well as significant IFNy secretion over no zipFv conditions (**Supplemental Fig 5C-D**).

### Characterization of differential binding affinity of CD33 and FLT3 zipFvs

Given the observed difference in dose-dependent activity of the FLT3 and CD33 zipFvs, we sought to investigate any potential competition or interference between them when both are present in the SUPRA system that could limit the efficacy against either target (**Fig 4A**). In this experiment, we created a dose-response matrix against each target as a co-culture of SUPRA CAR targeting single positive K562 targets with increasing amounts of “interfering” zipFv added into the system. From this assay, we see significant decreases in IFNy secretion when FLT3 zipFv is used to target FLT3+ K562 in the presence of increasing amounts of CD33 zipFv (**Fig 4B**). In contrast, the addition of FLT3 zipFv in the system targeting CD33+ K562 with CD33 zipFv does not show the same degree of decreased IFNy production, but we do observe a marked Hook effect at concentrations of CD33 zipFv above 25 ng (**Fig 4B**). Since the leucine zipper side of the zipFv protein is identical in both designs, this indicates that this behavior results from differences in the affinity of the FLT3 and CD33 scFvs to their cognate antigen.

**Figure 4.**
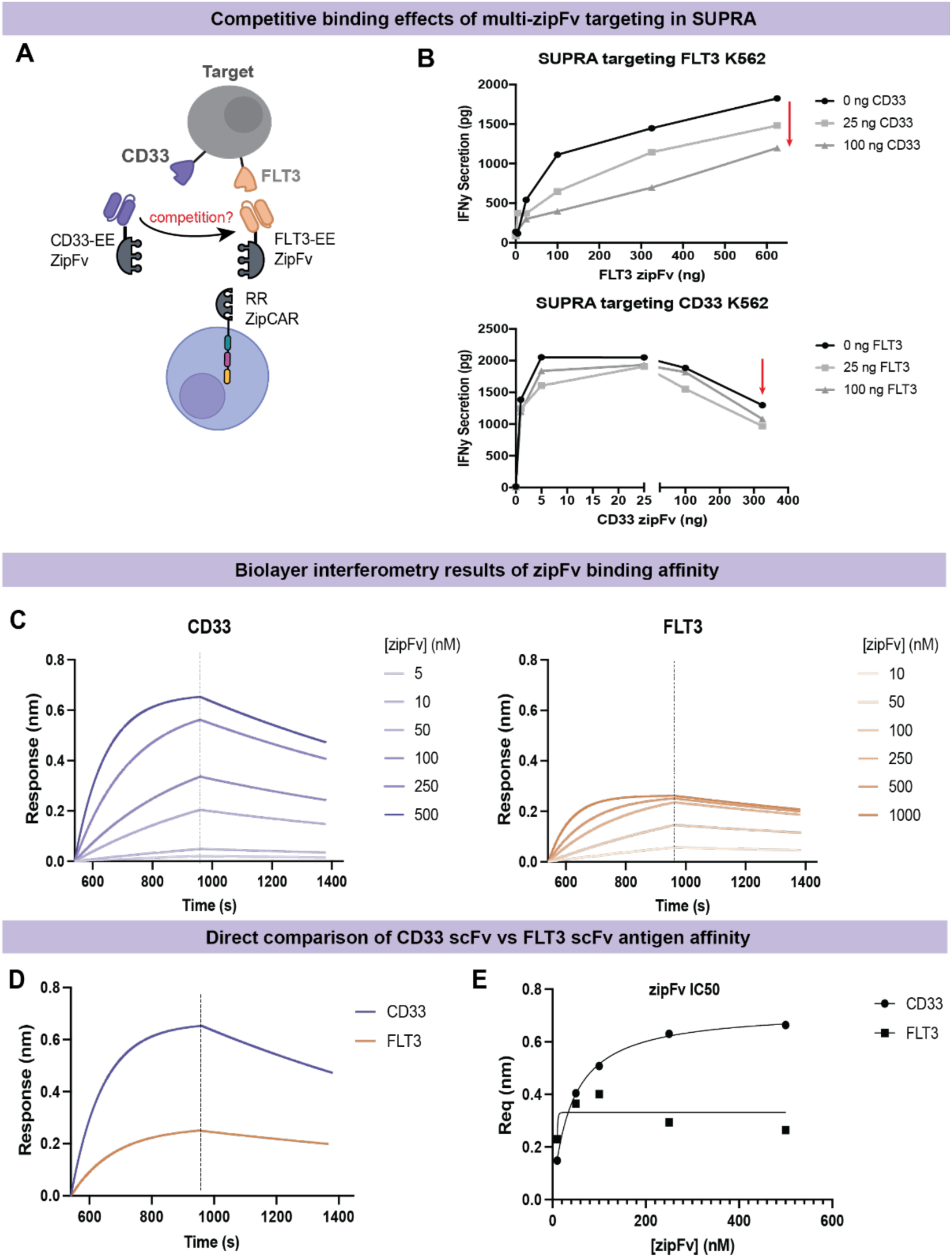
Characterization of competition due to differential affinity of FLT3 and CD33 scFvs to their cognate antigen. A. Schematic of potential competitive effect between zipFvs when both are present in the system. B. A dose competition assay was conducted by measuring IFNy secretion of the system when varying amounts of interfering zipFv were added to the co-culture. A decrease in activity against the FLT3+ K562 target is observed in the presence of increasing amounts of CD33 zipFv. This is not observed against CD33+ K562 targeting, although a Hook effect is observable at CD33 zipFv amounts over 25 ng. C. Biolayer interferometry of zipFv association and dissociation of tips coated in cognate antigen over a range of zipFv concentrations shows differential affinity of each zipFv. D. Direct comparison of BLI response at maximum zipFv dose shows a significantly higher association of CD33 zipFv compared to FLT3 zipFv. E. Plotting BLI response at equilibrium against the concentration of zipFv was used to calculate the IC50 of each protein, resulting in a CD33 IC_50_ of 41.0 nM and an FLT3 IC_50_ of 247.2 nM.

To further validate this difference, we performed bio-layer interferometry (BLI) assays to characterize the affinity of our zipFvs to FLT3 and CD33 antigens. Biotinylated CD33 and FLT3 proteins were immobilized onto streptavidin tips to characterize the association and dissociation of FLT3-NC7 and CD33-hu195 zipFv at various concentrations (**Fig 4C**). Based on the results, we clearly see a difference in affinity to their cognate antigen, with CD33 zipFv having a more rapid association (K_a_ = 1.81 × 10^4^ M^−1^s^−1^) compared to the FLT3 zipFv (K_a_ = 1.32 × 10^4^ M^−1^s^−1^) (**Fig 4D**). This difference is also highlighted by plotting BLI response at equilibrium against the concentration of zipFv to determine IC_50_ via non-linear regression, resulting in a CD33 IC_50_ of 41.0 nM and a FLT3 IC_50_ of 247.2 nM (**Fig 4E**). This difference in affinity can explain both the differential dose response and interference of CD33 zipFv on FLT3 targeting activity in our OR gate model, as well as highlight the importance of identifying optimal combinatorial dosing of all adapters in a split CAR system.

### *In vivo* validation of OR gate functionality in a xenograft mouse model

To test the OR gate *in vivo*, we first sought to establish a cancer cell line with a single positive target expression. The Nalm6 B-cell precursor ALL cell line was characterized to have moderate expression of CD33 antigen, and low expression of FLT3, designated as CD33^high^ Nalm6 (**Supplemental Fig 6A**). While Nalm6 is not an AML cell line, it is a well-characterized *in vivo* blood cancer model that is relatively easy to engineer to express our target antigens and enables initial *in vivo* validation of our system. To generate a FLT3+ line, Nalm6 cells were transduced to express human FLT3 antigen under antibiotic selection and designated as FLT3^high^ Nalm6 (**Supplemental Fig 6A**). We then validated that these cell lines can be effectively killed by our SUPRA system *in vitro* at the previously identified doses of 100 ng FLT3 zipFv and 25 ng CD33 zipFv in 24-hour co-culture cytotoxicity experiments at various E:T ratios as previously described (**Supplemental Fig 6B**). Lastly, to visualize both cell lines *in vivo* simultaneously, the FLT3^high^ Nalm6 cells were transduced to express Firefly luciferase while the CD33^high^ Nalm6 were transduced to express the orthogonal Antares luciferase (**Supplemental Fig 6C**).

We have shown previously that our SUPRA system has *in vivo* activity with zipFv dose ranges between 0.4 mg/kg and 5 mg/kg. Given the extensive dose range characterization we performed *in vitro*, we anticipated that some dose optimization would be needed *in vivo* as well, especially concerning the potential Hook effect with the high-affinity CD33 zipFv. To estimate the effective dose range, we calculated a rough scaling of the *in vitro* dose of 25 ng CD33 zipFv to an *in vivo* dose of 0.3 mg/kg given a total mouse blood volume of 2 mLs (**Supplemental Fig 7**). To cover a range across this dose and capture a high dose, we chose 0.15 mg/kg, 0.6 mg/kg, and 3 mg/kg of CD33 zipFv as an initial dosing trial against CD33^high^ Nalm6 with two mice per group following the dosing schedule shown in Figure 5A. Based on this initial trial, the 0.6 mg/kg and 0.15 mg/kg doses proved more effective at reducing tumor burden than the 3 mg/kg group and control groups containing zipCAR only or mock transduced T cells (**Supplemental Fig 8A**). For FLT3 zipFv dosing, we chose a higher range of 0.75 mg/kg, 1.5 mg/kg, and 3 mg/kg given its lower affinity than the CD33 zipFv. As expected, our initial dosing trials did not show a marked Hook effect for the FLT3 zipFv targeting FLT3^high^ Nalm, with the higher doses of 3 mg/kg and 1.5 mg/kg outperforming the lower dose of 0.75 mg/kg in controlling tumor burden (**Supplemental Fig 8B**). Given these initial trial results, we chose a dose of 0.6 mg/kg CD33 zipFv and 1.5 mg/kg FLT3 zipFv for our dual target model.

**Figure 5.**
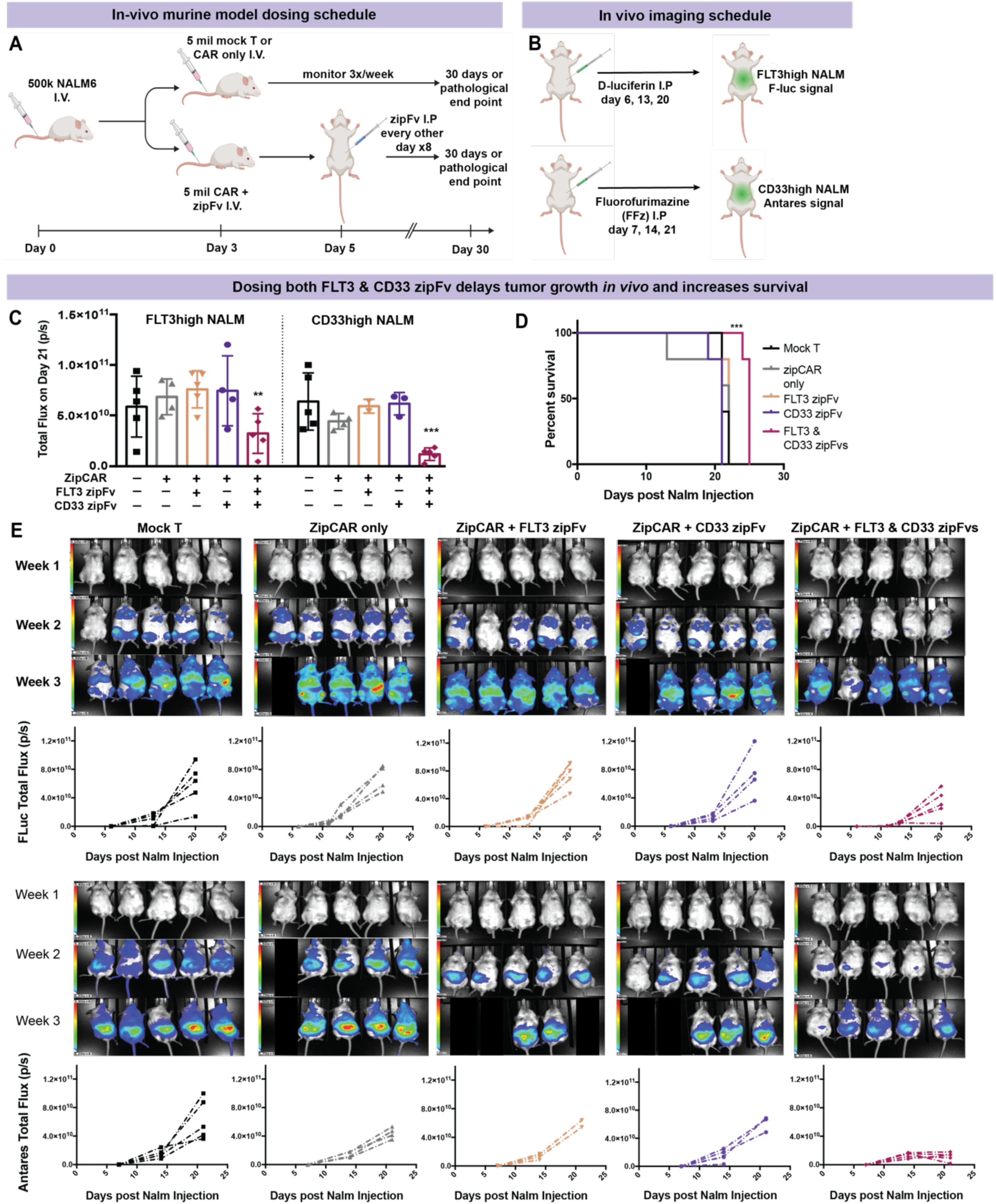
*In vivo* validation of SUPRA OR Gate in a xenograft mouse model. A. Dosing schedule of NOD SCID mice with Nalm6 cells followed by CAR T or mock transduced T cells along with corresponding zipFv. B. To independently image FLT3^high^ and CD33^high^ NALM tumor burden, the administration of D-luciferin and FFz were 24 hours apart to ensure residual substrate had cleared. C. Mice given both FLT3 and CD33 zipFv had significantly decreased tumor burden of both cell lines at week 3 compared to those with no zipFv or either zipFv alone. D. Mice given both FLT3 and CD33 zipFv had significantly increased survival rate than those given no zipFv or only FLT3 zipFv. E. Monitoring of FLT3^high^ Nalm6 growth via Firefly luciferase luminescence shows that FLT3 only zipFv administration was insufficient to control the tumor burden, while the group given both zipFvs had a significant reduction in tumor burden across the monitoring period leading to a substantial delay in tumor growth. Monitoring of CD33^high^ Nalm6 growth via Antares luciferase luminescence also shows that only the group given both CD33 and FLT3 zipFvs has a significant decrease in tumor growth rate across the treatment and sustained reduction in CD33^high^ Nalm6 cells.

To test the system against eliminating both CD33^high^ and FLT3^high^ Nalm6 cell lines simultaneously, each NOD SCID NSG mouse was injected i.v. with a 1:1 mix of both lines on Day 0 (250k each) followed by T cell injection of either zipCAR only, zipCAR incubated with the first dose of zipFv, or mock transduced CD3+ T cells on Day 3. Of the three groups that received the full SUPRA system, one group only received FLT3 zipFv, one group received only CD33 zipFv, while the complete OR gate group received both CD33 and FLT3 zipFvs and showed increased survival compared to the remaining groups (**Fig 5D**). Growth of the FLT3^high^ Nalm6-Firefly luciferase cells was monitored via IVIS imaging of the luciferase signal following i.p injection of D-Luciferin (**Fig 5B**), and while the FLT3 zipFv only group showed some delay in tumor growth as seen in the week 2 luminescence images, only the group given both zipFvs showed a significant reduction in FLT3^high^ tumor burden over the entire treatment period (**Fig 5C,E**). Growth of the CD33^high^ Nalm6-Antares luciferase cells was similarly monitored via IVIS imaging following the i.p injection of fluorofurimazine. Similarly, only the group given both FLT3 and CD33 zipFvs showed an effective reduction in tumor growth rate that persisted after the last dose was administered (**Fig 5C,E**). These studies indicate preliminary *in vivo* efficacy of dosing both zipFvs as an OR gate in a mixed target model and serve to highlight the importance of dose optimization in the development of effective therapies.

### HSPC sparing and identification of a potentially effective dose range

As a preliminary indication of the safety profile of our SUPRA system, we sought to identify a therapeutic window of activity *in-vitro* that spares primary human hematopoietic stem and progenitor cells (HSPCs) from ablation while retaining cytotoxic activity against cancer cell lines. We obtained bone marrow or cord blood derived CD34+ enriched human HSPCs (StemCell) and characterized their expression of the target antigens CD33 and FLT3 using the same antibody binding capacity assay previously described (**Fig 6A**). The bulk CD34+ population of both stem and progenitor cells showed moderate CD33 expression and low FLT3 expression (**Fig 6B**). CD90 has been identified as a marker that differentiates HSCs from multipotent, erythro-myeloid, and lympho-myeloid progenitors^46^. By gating for the CD34+CD90+ subset, we quantified the expression of CD33 and FLT3 within this population and found it to be slightly enriched in FLT3 and lower in CD33 expression (**Fig 6A,C**). Given the limited number of this HSC-enriched population, we chose one E:T ratio of 2.5:1 to test the dose-dependent cytotoxic activity of the SUPRA system against these cells in a 24-hour co-culture. To assess cytotoxicity, HSPCs were isolated from T cells as CD3-CD34+ followed by further gating of a CD90+CD45RA-HSC-enriched subset (**Fig 6D**). Compared to the cytotoxic activity against our in-vivo model NALM cell lines, we observed minimal toxicity when targeting FLT3 that was slightly higher in the HSC-enriched subset, and a maximum of ∼30% toxicity when targeting CD33 at our highest dosage (**Fig 6E**). Given our utilized therapeutic dosages of 100-125 ng of FLT3 zipFv and 25 ng of CD33 zipFv, there is minimal toxicity against the HSPCs at these doses, indicating the existence of a therapeutic dose window for effective specific lysis of cancer cells with minimal off-tumor activity.

**Figure 6.**
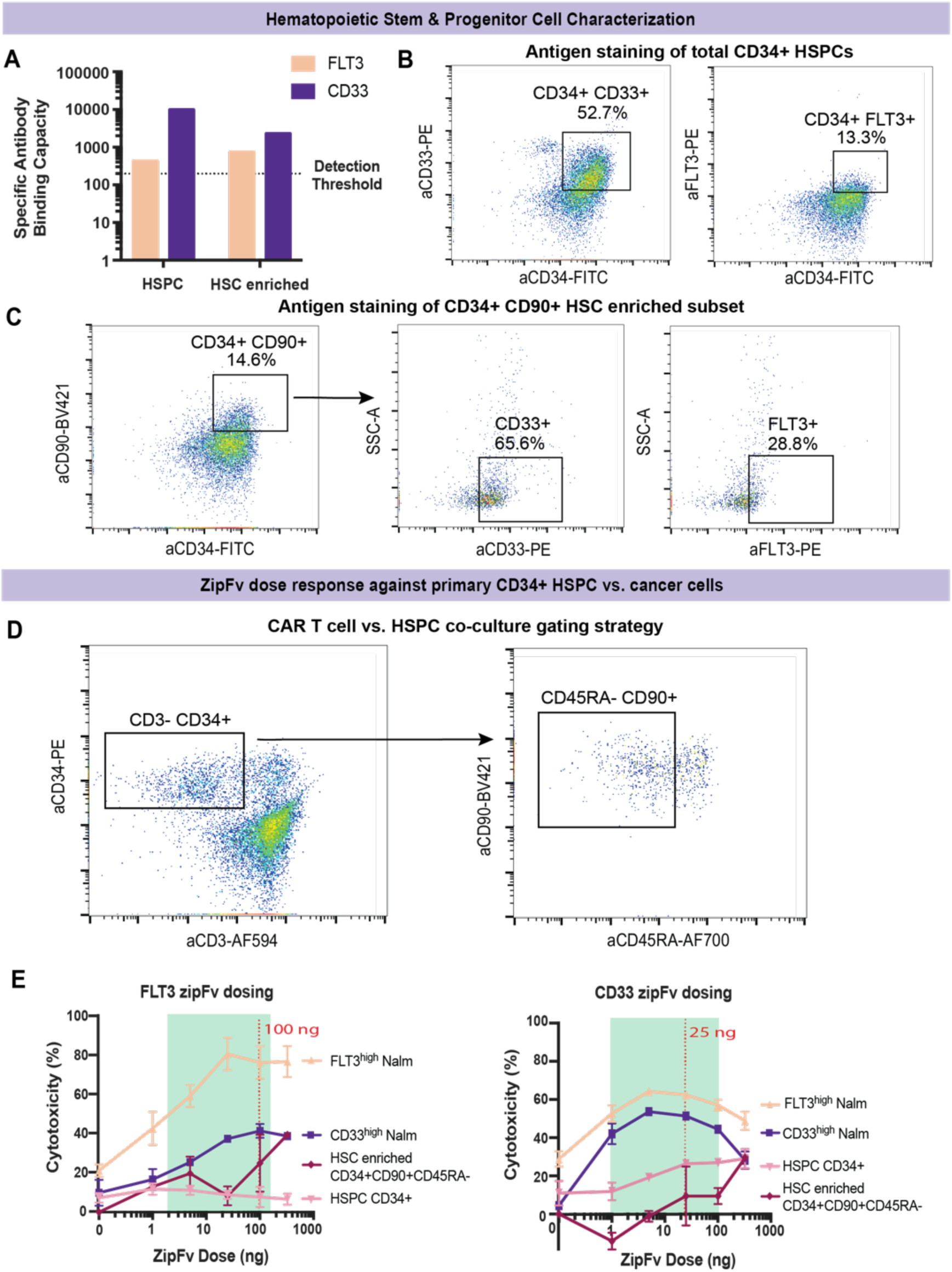
Identification of a therapeutic zipFv dose window that spares healthy HSPCs *in vitro*. A. CD34 enriched HSPCs were obtained as cryopreserved vials from StemCell and stained for antigen expression. MESF quantification of total CD34+ HSPC expression of the target antigens shows a moderate expression of CD33 and low expression of FLT3 while the CD90+ stem-like subset had lower CD33 and higher FLT3 expression. B. HSPCs were gated for CD34 expression along with either antigen of interest. C. The subpopulation of CD90+ cells makes up 10-15% of the total HSPC population with varying levels of antigen expression. D. In co-culture conditions with the SUPRA system, HSPCs were first gated as CD3-, CD34+ followed by further subclassification of an HSC-enriched subset as CD45RA-CD90+ within the HSPC population. E. Primary CD3+ T cells expressing RR zipCAR were co-cultured with healthy HSPCs at an E:T of 2.5:1 for 24 hours in the presence of a range of zipFv doses. By comparing to the cytotoxic activity against our Nalm6 target cells, we can identify a therapeutic window of >50% activity against the target cells that has <30% toxicity against the HSPCs for both zipFvs.

## Discussion

CAR T cell therapies have proven to be a remarkable breakthrough and exciting area of progress in immunotherapy with long-term proven efficacy^47^. Currently implemented CAR designs suffer from fixed antigen targeting capabilities, allowing for antigen escape, as well as high levels of on-target, off-tumor toxicity due to their non-tunability^48–53^. To overcome these limitations, we have previously introduced our SUPRA CAR platform as a tunable, multiplexed targeting platform. In this study, we adapted SUPRA towards treating AML, a heterogeneous disease with a very poor prognosis that has proven difficult to address with conventional CAR T designs. Among the many antigen markers that have been characterized for AML, we focused on CD33, a myeloid marker shown to be expressed on over 85% of AML cases, and FLT3, a common mutation in AML that has been tied to poor disease progression. We show that we can dose two zipFv adapters simultaneously to achieve combinatorial SUPRA targeting that eliminates target AML cell lines with varying expression levels of CD33 and FLT3. Additionally, we characterized the potential competition present in this system to identify concentrations of each adapter that limit inhibition and account for the variable affinity of each scFv to their target antigen. Interestingly, this translated to in-vivo dose optimization of FLT3 and CD33 zipFvs, further emphasizing the importance of adapter characterization in split systems. Within our working dose range, we confirmed minimal toxicity against bone marrow or cord blood derived CD34+ HSPCs as well as the smaller CD90+CD45RA-HSC-enriched subset, demonstrating the potential of SUPRA as a treatment strategy that overcomes the high toxicity of conventional CAR treatment.

In our *in-vitro* screening experiments, we consistently saw lower basal activation of the zipCAR compared to conventional CAR designs. This implies that SUPRA has a higher activation threshold, making it less sensitive but more specific to our target antigens. While this addresses safety concerns of off-target activity, it also raises a question of potency of the therapy in effectively eliminating all cancer cells. Due to the tunable nature of this design, dosing can be adjusted to compensate for this lower sensitivity if a stronger response is needed. Indeed, we can achieve higher cytokine secretion at optimal dosages compared to conventional CAR counterparts. In early 2020, AbbVie and Calibr initiated the first clinical trial of a “switchable” CAR design composed of an antibody switch as the adapter to the CAR, motivated by the need to develop safer and more controllable CAR designs^54^. Initial results shared in late 2022 showed that 6 out of 9 patients treated had a complete response with only one dose CAR-T and cognate antibody switch, SWI019^55^. Additionally, these patients experienced lower levels of CRS toxicity, if at all. These preliminary results are an exciting demonstration of the potential of split CAR designs, such as SUPRA, to combat conventional CAR T treatment toxicity while retaining therapeutic efficacy.

In this study, we focused on just two target antigens, CD33 and FLT3, to demonstrate the feasibility of a SUPRA OR gate against AML. However, the flexibility of the SUPRA system offers the potential to switch or add additional antigen targets depending on the individual patient’s disease phenotype. We have also demonstrated both “AND” and “NOT” logic in SUPRA previously, which can be coupled with this OR gate design to further increase the specificity of the circuit against cancer target cells. Given the characterization we have done here, it will be important to explore the interaction between adapter binding affinity and the relative abundance of each target to design effective combinatorial targeting strategies as we increase the complexity of these systems.

## ACKNOWLEDGEMENTS

W.W.W. acknowledges funding from NIH Award (U01CA265713). M.Y.S. acknowledges funding from NIH F31 Award (1F31HL162477-01A1). We also thank Wong lab members for suggestions on the manuscript; Grinstaff lab members for assistance on the BLI assay; BU IVIS imaging core facility for mice *in vivo* imaging (NIH grant 1S10RR024523-01).

## AUTHOR CONTRIBUTIONS

M.Y.S designed and performed primary T cell and HSPC experiments, analyzed the data, and generated figures. J.C. designed and performed Jurkat screening experiments, and assisted M.Y.S with primary T cell experiments and *in vivo* studies. M.L. performed BLI assays and generated corresponding figures. S.L. assisted with *in vitro* zipFv functional screening. H.D., Y.L., N.L., and T.L designed and generated CD33 and FLT3 binders and zipFv constructs, engineered K562 target cells lines, and advised on *in vivo* and HSPC experiments with input from N.F, B.S.G., and M.G.A. W.W.W supervised the project and analyzed the data. All authors commented on and approved the paper.

## DECLARATION OF INTERESTS

W.W.W. is a scientific co-founder and shareholder of Senti Biosciences and 4Immune.

## Methods

### Cell line culture

HEK293T cells were cultured in complete DMEM (Corning) containing 10% FBS (Thermo Fisher), 100 units/mL penicillin (Corning), 100 ug/mL streptomycin (Corning), 1% L-glutamine (Corning), and 1 mM sodium pyruvate (Lonza). Jurkat T cells were cultured in complete RPMI (Cytvia) containing 5% FBS, 100 units/mL penicillin, 100 ug/mL streptomycin, and 1% L-glutamine. K562 cells, MV4-11 cells, THP-1 cells, and NALM cells were all cultured in complete RPMI containing 10% FBS, 100 units/mL penicillin, 100 ug/mL streptomycin, and 1% L-glutamine.

### zipFv and zipCAR construct design

zipFvs were designed by Senti Biosciences as fusions of scFvs from known antibody clones (FLT3 clones NC-7 and D43, CD33 clones Hu195 and Mylotarg) to EE or JUN leucine zippers via a GC linker, followed by a 6X His tag for purification (Figure S1).

zipCARs were designed as previously described in the original SUPRA system as third generation CAR constructs expressed under an SFFV promoter in a lentiviral backbone (Figure S1). For the initial zipFv screening, both RR and JUN zipCARs were evaluated along with their cognate zipFv pairs. All zipCARs contained a myc tag c-terminal to the leucine zipper and were fused with mCherry after CD3z to assess expression levels.

### Expression and purification of zipFv

For zipFv protein production, Freestyle 293-F cells (Thermo Scientific) were transfected with pSecTag2A mammalian expression vector based on the supplier’s protocol. Three days post transfection, zipFv was collected from the cell supernatant through centrifugation at 400 x g for 5 minutes. The collected zipFvs were then purified by the Probond purification system with native conditions.

### Western Blot and SDS-PAGE Gel Electrophoresis

To verify the expression of zipFv proteins, western blot was performed based on standard protocol. Purified proteins were first mixed with NuPAGE LDS sample buffer (Fisher Scientific) and NuPAGE reducing agent (Fisher Scientific), and then heated on a thermocycler at 90°C for 10 minutes. Denatured samples were then loaded on SDS-PAGE gel in NuPAGE MOPS SDS running buffer (Thermo Scientific) and run at 200V for 30 minutes. iBlot 2 gel transfer device was used to transfer protein to the membrane, which was then blocked by 5% of milk powder (Research Products International) in 1X PBS-Tween 20 (Thermo Scientific). To detect produced zipFvs, the blocked membrane was incubated overnight with 1:1000 diluted 6x-His Tag Monoclonal Antibody (4E3D10H2/E3) in PBS-Tween 20. Images were captured the next day using Gel Doc EZ imager (Biorad).

### Primary human T cell isolation and culture

Primary CD3+ T cells were isolated from anonymized whole peripheral blood obtained from Boston Children’s Hospital using the STEMCELL RosetteSep Human T Cell Enrichment Cocktail. After isolation, CD3+ T cells were cryopreserved in 90% human Ab serum (Innovative Research) and 10% DMSO. For cell culture, complete T cell media consisted of X-VIVO 15 (Lonza) supplemented with 5% human Ab serum and 55 µM 2-mercaptoethanol. 50 units/mL of IL-2 (NCI BRB Preclinical Repository) was added to the media upon addition to the T cell culture.

### Lentivirus generation and transduction of human T cells

Replication-incompetent lentivirus was packaged using HEK293T Lenti-X cells (Takara) and third generation lentiviral packaging and envelope plasmids. Briefly, HEK293T cells plated in a 125 mm^2^ culture plate were PEI transfected with the lentiviral and zipCAR plasmids at 80-90% confluency. 24 hours post transfection, HEK293T culture media was removed and replaced with FreeStyle 293 expression medium (Gibco) supplemented with 100 µg/mL streptomycin, 100 units/mL penicillin, 1 mM sodium pyruvate, and 5 mM sodium butyrate. Lentivirus-containing supernatant was collected two times between 48 and 96 hours post transfection and replaced with fresh complete FreeStyle media between collections. The harvested lentivirus was concentrated using a 40% w/v PEG-8000 and 1.2 M NaCl solution overnight, then spun down for 1 hr at 1600xg at 4°C. The supernatant was then poured off, and the viral pellet was resuspended in 4 mL cold PBS. Concentrated virus was either used immediately or flash frozen in liquid nitrogen and stored at −80°C.

T cells were thawed 48-72 hrs before transduction and cultured in complete X-VIVO 15 media for 24 hrs, then activated with Immunocult human CD3/CD28 T cell activator (STEMCELL) for up to 48 hrs. One day before transduction, a non-TC treated 6 or 12-well plate was coated with 20 ug/mL Retronectin (Takara) in PBS and incubated at 4°C overnight. The Retronectin solution was then removed, and the plate was blocked with 2% BSA solution for 30 minutes before adding 2 mL or 1 mL of concentrated virus per well for either a 6-well or 12-well transduction, respectively. This was spun down at 1200xg for 90 minutes, then removed and activated primary CD3+ T cells were added at a concentration of 250k cells/mL and incubated at 37°C. Transduction efficiency was assessed as mCherry expression via flow cytometry, and transduced T cells were maintained between 500k and 1 mil cells/mL for 10-14 days for experimental use.

### *In-vitro* NFAT Jurkat activation assay

On the day of transduction, 2 mL NFAT Jurkat cells were mixed with 50 uL of concentrated virus per well for a 6-well plate at a concentration of 250k cells/mL. The plate was spun down at 1000xg for 30 minutes then the virus containing media was replaced with fresh Jurkat culture media by spinning down at 400xg for 5 minutes. Transduction efficiency was assessed by mCherry expression through flow cytometry 3 days after transduction. Transduced NFAT Jurkat cells were maintained between 250k and 1.5 mil cells/mL.

NFAT Jurkat activation assay was performed by incubating transduced NFAT-GFP Jurkat cells expressing zipCAR with engineered K562 target cells at an E:T ratio of 1:1 (100k NFAT Jurkat cells and 100k target cells) with 25 ng/well of each zipFv for 16-20 hours at 37°C. Activation of NFAT Jurkat cells was assessed by NFAT-GFP expression through flow cytometry.

### *In-vitro* cytotoxicity assay

Co-culture cytotoxicity assays were initially performed by incubating primary human CD3+ T cells expressing zipCAR with engineered K562 target cells at an E:T ratio of 2:1 (50k CAR T cells and 25k target cells) along with a range of zipFv doses from 0-625 ng/well for 24 hours at 37°C. Based on these initial assays, a dose of 125 ng of FLT3 zipFv and 25 ng of CD33 zipFv was used for subsequent assays that varied E:T ratio against K562, MV4-11, and THP-1 cell lines. For all E:T variations, a target cell number of 20k was used with CAR T amounts varying from 20k for a 1:1 ratio to 200k for a 10:1 ratio. Target cells were stained with either eFluor 450 or CellTrace Violet prior to co-culture, and after 24 hours, the supernatant was saved, and the cells were resuspended in FACS buffer to analyze via flow cytometry. Cytotoxicity was measured as viable target cell count compared to target cell only conditions by gating for eFluor 450+ or CellTrace Violet+ cells. In the mixed single positive K562 co-culture assay, CD33+ K562 cells express GFP, allowing for specific gating based on the GFP signal.

### Cytokine release assay

Supernatant saved from cytotoxicity assays was analyzed using an enzyme-linked immunosorbent assay (ELISA) to measure IFNγ secretion. The BD OptEIA Human IFNγ ELISA kit was used along with BD Reagent Set B according to the manufacturer’s instructions. Supernatant was diluted at either a 1:50 or 1:100 ratio in assay diluent to ensure cytokine amounts were within the detection limits of the kit.

### zipFv competition assay

To evaluate the potential competition between FLT3 and CD33 zipFvs when dosed simultaneously, co-culture assays were performed as previously described at an E:T ratio of 1:1 in a dose response matrix. Briefly, CAR T cells were cultured with one target cell type, either WT K562, FLT3^high^ K562, or CD33^high^ K562, along with varying doses of “targeting” and “non-targeting” zipFv from 0-625 ng/well such that at each dose of targeting zipFv, a range of non-targeting zipFv doses was added to evaluate interference. After 24 hours of co-culture at 37°C, the supernatant was saved and analyzed via ELISA to measure IFNγ secretion as previously described.

### Biolayer interferometry assay

CD33-biotin was purchased from Sino Biological, and FLT3-His was purchased (Sino Biological) and then biotinylated in-house using the EZ-Link Biotinylation kit (Fisher Scientific). After equilibrating tips in BLI buffer (1X PBS + 0.05% Tween-20) and establishing a baseline, the streptavidin (SA) coated biosensor tips were dipped into wells containing 10 µg/mL of CD33 or FLT3 (separately) for immobilization, followed by transfer into the buffer to establish baseline 2. Loaded tips were then dipped into various concentrations of zipFv (0-1000 nM) to allow for association before transfer back into the buffer for dissociation. Analysis was performed in Octet Analysis Studio 13.0. Association was controlled by subtraction of a trace where the tip was not loaded with protein before exposure to zipFv, and dissociation was controlled via subtraction of a trace containing loaded protein but no exposure to zipFv. Y-axis was aligned to the last 5 seconds of the second baseline, and data was fit to a Mass Transport model.

### HSPC sparing assay

Bone marrow or cord blood derived human primary CD34 enriched cells were obtained from StemCell and cultured in StemSpan SFEM II Human Hematopoietic Stem Cell Culture Medium (StemCell). Cells were thawed and stained for antigen expression then used in co-culture cytotoxicity assays within 24 hours as described previously, with an E:T ratio of 2:1 and varying concentrations of each targeting zipFv. After 24 hours of co-culture, the supernatant was saved, and cells were resuspended in FACS buffer for analysis via flow cytometry. Cytotoxicity was measured as viable CD3-CD34+ HSPC cell count compared to target cell only and zero zipFv conditions. HSC-enriched subset cytotoxicity was measured by further gating the HSPC population on CD90+CD45RA-.

### Xenograft mouse model

Cohorts of female NOD.Cg-Prkdcscid Il2rgtm1Wjl/SzJ (NSG) mice were ordered from the Jackson Laboratories and housed in the BUMC Animal Science Center. All animal protocols were reviewed and approved by the Institutional Animal Care and Use Committee (IACUC) at Boston University. Briefly, NSG mice were injected with 500k luciferase-expressing NALM6 cells intravenously on day 0. CD33^high^ NALM6 were engineered to express Antares luciferase while FLT3^high^ NALM6 were engineered to express Firefly luciferase for a dual reporter system. In animals containing both target cells, they were mixed at 250k each for a total cancer cell injection of 500k/mouse. On day 3 or day 6, 5 million CAR-T cells were injected intravenously. For conditions containing zipFv, CAR T cells were incubated with the first dose zipFv 30 minutes prior to injection. Subsequently, zipFv was injected intraperitoneally every other day for a total of 4 or 6 injections. Tumor burden was measured by IVIS Spectrum and was quantified as total flux (photons/sec) in the entire mouse. Images were acquired within 10 minutes of intraperitoneal injection of 150 mg/kg of D-luciferin (Perkin Elmer) or 17.6 nM of FFz (Promega). Images were analyzed using Living Image (Perkin Elmer).

### Statistical analysis

To evaluate statistical significance, data between groups were evaluated using an unpaired two-tailed t test. P values are reported (not significant = p > 0.05, ∗ = p < 0.05, ∗∗ = p < 0.01, ∗∗∗ = p < 0.001), and error bars are plotted as standard deviations between replicates.

## RESOURCE AVAILABILITY

### Lead contact

Further information and requests for resources and reagents should be directed to and will be fulfilled by the Lead Contact, Wilson Wong.

### Materials availability

All plasmid constructs and cell lines generated in this study will be made available on request, but we may require a payment and/or a completed Materials Transfer Agreement if there is potential for commercial application.

### Data and core availability

All data reported in this paper will be shared by the lead contact upon request. This paper does not report original code. Any additional information required to reanalyze the data reported in this paper is available from the lead contact upon request.

## References

1. Döhner, H., Weisdorf, D. J. & Bloomfield, C. D. Acute Myeloid Leukemia. 10.1056/NEJMra1406184 373, 1136–1152 (2015).

2. Acute Myeloid Leukemia — Cancer Stat Facts. https://seer.cancer.gov/statfacts/html/amyl.html.

3. Long, L. et al. Genetic biomarkers of drug resistance: A compass of prognosis and targeted therapy in acute myeloid leukemia. Drug Resistance Updates 52, 100703 (2020).

4. Bazarbachi, A. et al. Clinical practice recommendation on hematopoietic stem cell transplantation for acute myeloid leukemia patients with FLT3-internal tandem duplication: a position statement from the Acute Leukemia Working Party of the European Society for Blood and Marrow Transplantation. Haematologica 105, 1507 (2020).

5. Cornelissen, J. J. & Blaise, D. Hematopoietic stem cell transplantation for patients with AML in first complete remission. Blood 127, 62–70 (2016).

6. Takami, A. Hematopoietic stem cell transplantation for acute myeloid leukemia. Int J Hematol 107, 513–518 (2018).

7. Short, N. J. et al. Advances in the treatment of acute myeloid leukemia: New drugs and new challenges. Cancer Discov 10, 506–525 (2020).

8. Capria, S. et al. Autologous stem cell transplantation in favorable-risk acute myeloid leukemia: single-center experience and current challenges. Int J Hematol 116, 586–593 (2022).

9. Awada, H. et al. A Focus on Intermediate-Risk Acute Myeloid Leukemia: Sub-Classification Updates and Therapeutic Challenges. Cancers 2022, Vol. 14, Page 4166 14, 4166 (2022).

10. Stelmach, P. & Trumpp, A. Leukemic stem cells and therapy resistance in acute myeloid leukemia. Haematologica 108, 353 (2023).

11. C. King, PharmD, BCOP, A. & S. Orozco, PharmD, J. Axicabtagene Ciloleucel: The First FDA-Approved CAR T-Cell Therapy for Relapsed/Refractory Large B-Cell Lymphoma. J Adv Pract Oncol 10, 878 (2019).

12. Maude, S. L. et al. Tisagenlecleucel in Children and Young Adults with B-Cell Lymphoblastic Leukemia. N Engl J Med 378, 439 (2018).

13. Abramson, J. S. et al. Lisocabtagene maraleucel for patients with relapsed or refractory large B-cell lymphomas (TRANSCEND NHL 001): a multicentre seamless design study. The Lancet 396, 839–852 (2020).

14. Munshi, N. C. et al. Idecabtagene Vicleucel in Relapsed and Refractory Multiple Myeloma. New England Journal of Medicine 384, 705–716 (2021).

15. Wang, Y. et al. Brexucabtagene Autoleucel for Relapsed/Refractory Mantle Cell Lymphoma: Real World Experience from the US Lymphoma CAR T Consortium. Blood 138, 744 (2021).

16. Berdeja, J. G. et al. Ciltacabtagene autoleucel, a B-cell maturation antigen-directed chimeric antigen receptor T-cell therapy in patients with relapsed or refractory multiple myeloma (CARTITUDE-1): a phase 1b/2 open-label study. The Lancet 398, 314–324 (2021).

17. Zhao, J., Song, Y. & Liu, D. Clinical trials of dual-target CAR T cells, donor-derived CAR T cells, and universal CAR T cells for acute lymphoid leukemia. Journal of Hematology & Oncology 2019 12:1 12, 1–11 (2019).

18. Mardiana, S. & Gill, S. CAR T Cells for Acute Myeloid Leukemia: State of the Art and Future Directions. Front Oncol 10, 697 (2020).

19. Zeng, A. G. X. et al. A cellular hierarchy framework for understanding heterogeneity and predicting drug response in acute myeloid leukemia. Nature Medicine 2022 28:6 28, 1212–1223 (2022).

20. Romer-Seibert, J. S. & Meyer, S. E. Genetic Heterogeneity and Clonal Evolution in Acute Myeloid Leukemia. Curr Opin Hematol 28, 64 (2021).

21. Horibata, S. et al. Heterogeneity in refractory acute myeloid leukemia. Proc Natl Acad Sci U S A 116, 10494–10503 (2019).

22. Schorr, C. & Perna, F. Targets for chimeric antigen receptor T-cell therapy of acute myeloid leukemia. Front Immunol 13, (2022).

23. Wang, Q. S. et al. Treatment of CD33-directed chimeric antigen receptor-modified T cells in one patient with relapsed and refractory acute myeloid leukemia. Molecular Therapy 23, 184–191 (2015).

24. Cho, J. H. et al. Engineering advanced logic and distributed computing in human CAR immune cells. Nature Communications 2021 12:1 12, 1–14 (2021).

25. Moghimi, B. et al. Preclinical assessment of the efficacy and specificity of GD2-B7H3 SynNotch CAR-T in metastatic neuroblastoma. Nature Communications 2021 12:1 12, 1–15 (2021).

26. Fedorov, V. D., Themeli, M. & Sadelain, M. PD-1– and CTLA-4–Based Inhibitory Chimeric Antigen Receptors (iCARs) Divert Off-Target Immunotherapy Responses. Sci Transl Med 5, 215ra172 (2013).

27. Zah, E. et al. Systematically optimized BCMA/CS1 bispecific CAR-T cells robustly control heterogeneous multiple myeloma. Nature Communications 2020 11:1 11, 1–13 (2020).

28. Srivastava, S. et al. Logic-Gated ROR1 Chimeric Antigen Receptor Expression Rescues T Cell-Mediated Toxicity to Normal Tissues and Enables Selective Tumor Targeting. Cancer Cell 35, 489–503.e8 (2019).

29. Roybal, K. T. et al. Precision Tumor Recognition by T Cells With Combinatorial Antigen-Sensing Circuits. Cell 164, 770–779 (2016).

30. Salzer, B. et al. Engineering AvidCARs for combinatorial antigen recognition and reversible control of CAR function. Nature Communications 2020 11:1 11, 1–16 (2020).

31. Lanitis, E., Coukos, G. & Irving, M. All systems go: converging synthetic biology and combinatorial treatment for CAR-T cell therapy. Curr Opin Biotechnol 65, 75–87 (2020).

32. Savanur, M. A., Weinstein-Marom, H. & Gross, G. Implementing Logic Gates for Safer Immunotherapy of Cancer. Front Immunol 12, 4678 (2021).

33. Haubner, S. et al. Cooperative CAR targeting to selectively eliminate AML and minimize escape. Cancer Cell (2023) doi:10.1016/J.CCELL.2023.09.010.

34. Wermke, M. et al. Proof of concept for a rapidly switchable universal CAR-T platform with UniCAR-T-CD123 in relapsed/refractory AML. Blood 137, 3145–3148 (2021).

35. Cho, J. H., Collins, J. J. & Wong, W. W. Universal Chimeric Antigen Receptors for Multiplexed and Logical Control of T Cell Responses. Cell 173, 1426–1438.e11 (2018).

36. Niswander, L. M. et al. Potent preclinical activity of FLT3-directed chimeric antigen receptor T-cell immunotherapy against *FLT3*-mutant acute myeloid leukemia and *KMT2A*-rearranged acute lymphoblastic leukemia. Haematologica 108, 457–471 (2023).

37. Wang, Y. et al. Targeting FLT3 in acute myeloid leukemia using ligand-based chimeric antigen receptor-engineered T cells. J Hematol Oncol 11, 1–12 (2018).

38. Godwin, C. D., Gale, R. P. & Walter, R. B. Gemtuzumab ozogamicin in acute myeloid leukemia. Leukemia 2017 31:9 31, 1855–1868 (2017).

39. Molica, M. et al. CD33 Expression and Gentuzumab Ozogamicin in Acute Myeloid Leukemia: Two Sides of the Same Coin. Cancers (Basel) 13, (2021).

40. Cortes, J. E. et al. Prevention, recognition, and management of adverse events associated with gemtuzumab ozogamicin use in acute myeloid leukemia. J Hematol Oncol 13, 1–8 (2020).

41. Subklewe, M. et al. Preliminary Results from a Phase 1 First-in-Human Study of AMG 673, a Novel Half-Life Extended (HLE) Anti-CD33/CD3 BiTE® (Bispecific T-Cell Engager) in Patients with Relapsed/Refractory (R/R) Acute Myeloid Leukemia (AML). Blood 134, 833 (2019).

42. Marcinek, A. et al. CD33 BiTE® molecule-mediated immune synapse formation and subsequent T-cell activation is determined by the expression profile of activating and inhibitory checkpoint molecules on AML cells. Cancer Immunology, Immunotherapy 1–14 (2023) doi:10.1007/S00262-023-03439-X/FIGURES/6.

43. Daver, N., Schlenk, R. F., Russell, N. H. & Levis, M. J. Targeting FLT3 mutations in AML: review of current knowledge and evidence. Leukemia 2019 33:2 33, 299–312 (2019).

44. Giannakopoulou, E. et al. A T cell receptor targeting a recurrent driver mutation in FLT3 mediates elimination of primary human acute myeloid leukemia in vivo. Nature Cancer 2023 1–17 (2023) doi:10.1038/s43018-023-00642-8.

45. Marofi, F. et al. Novel CAR T therapy is a ray of hope in the treatment of seriously ill AML patients. Stem Cell Research & Therapy 2021 12:1 12, 1–23 (2021).

46. Radtke, S. et al. Isolation of a Highly Purified HSC-enriched CD34+CD90+CD45RA− Cell Subset for Allogeneic Transplantation in the Nonhuman Primate Large-animal Model. Transplant Direct 6, (2020).

47. Melenhorst, J. J. et al. Decade-long leukaemia remissions with persistence of CD4+ CAR T cells. Nature 2022 602:7897 602, 503–509 (2022).

48. Zhang, Y. et al. Single-Cell Analysis of Target Antigens of CAR-T Reveals a Potential Landscape of “On-Target, Off-Tumor Toxicity”. Front Immunol 12, 799206 (2021).

49. Flugel, C. L. et al. Overcoming on-target, off-tumour toxicity of CAR T cell therapy for solid tumours. Nature Reviews Clinical Oncology 2022 20:1 20, 49–62 (2022).

50. Santomasso, B. D. et al. Clinical and Biologic Correlates of Neurotoxicity Associated with CAR T Cell Therapy in Patients with B-cell Acute Lymphoblastic Leukemia (B-ALL). Cancer Discov 8, 958 (2018).

51. Frey, N. V. & Porter, D. L. Cytokine release syndrome with novel therapeutics for acute lymphoblastic leukemia. Hematology 2016, 567–572 (2016).

52. Neelapu, S. S. et al. Chimeric antigen receptor T-cell therapy - assessment and management of toxicities. Nat Rev Clin Oncol 15, 47–62 (2018).

53. Sterner, R. C. & Sterner, R. M. CAR-T cell therapy: current limitations and potential strategies. Blood Cancer Journal 2021 11:4 11, 1–11 (2021).

54. AbbVie & Scripps-based Calibr Moves Novel ‘Switchable’ CAR-T Technology to Phase I Clinical Trial. https://www.trialsitenews.com/a/abbvie-scripps-based-calibr-moves-novel-switchable-car-t-technology-to-phase-i-clinical-trial.

55. Calibr reports promising results from first-in-human clinical trial of switchable CAR-T (CLBR001 + SWI019), a next-generation universal CAR-T platform designed to enhance the versatility and safety of cell therapies | Scripps Research. https://www.scripps.edu/news-and-events/press-room/2022/20220921-calibr-cart.html.

