## Supplemental Figures for "Tunable Universal OR-gated CAR T cells for AML"

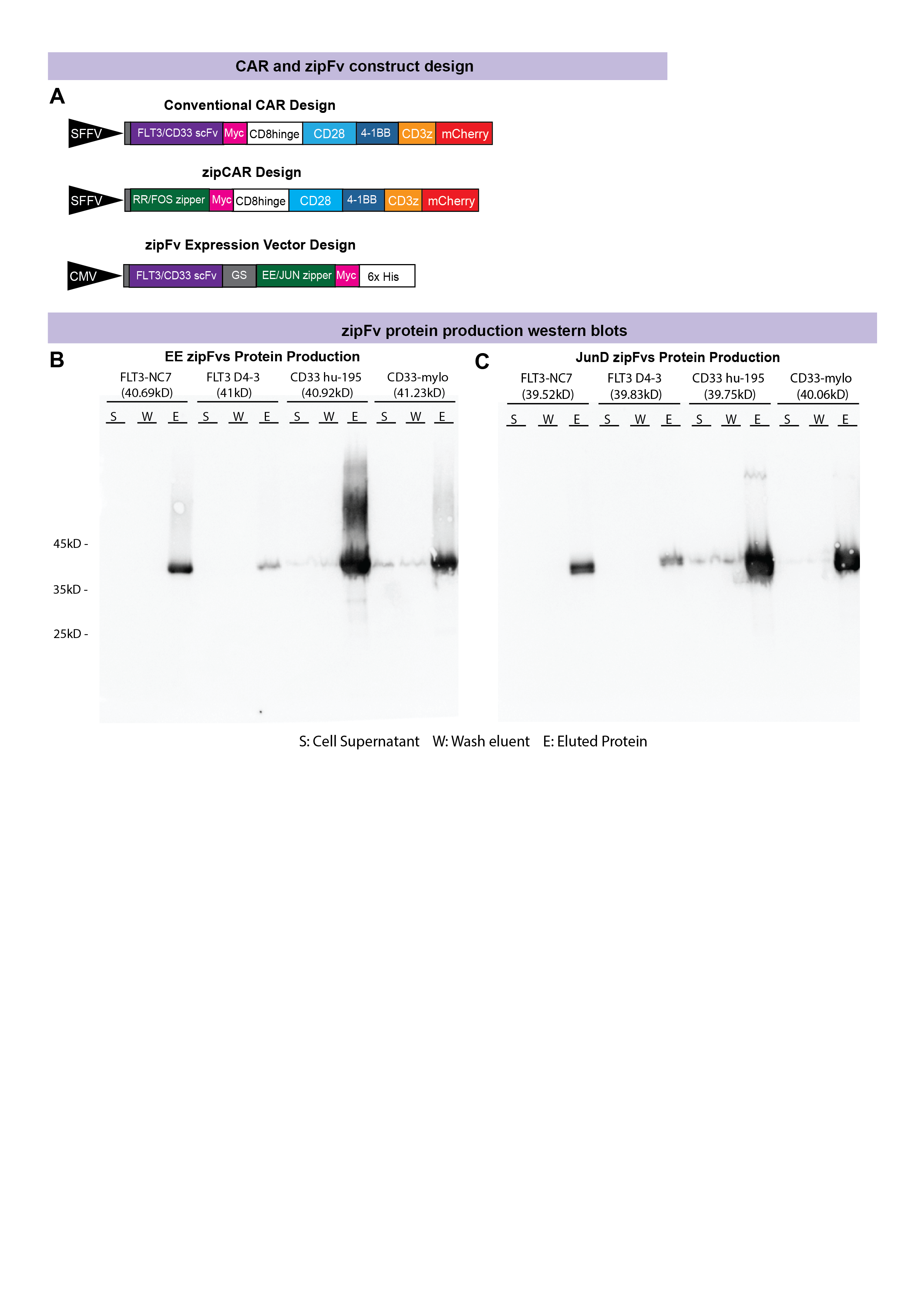

**Supplemental Figure 1. SUPRA component design and zipFv protein production.** A. CAR constructs were designed as 3^rd^ generation CARs with either FLT3 scFv, CD33 scFv, or RR/FOS leucine zipper halves followed by a myc tag, CD8 hinge and transmembrane domain, CD28 and 4-1BB costimulatory domains, the CD3z activation domain and an mCherry for visualization of expression. ZipFv expression vector was designed as the FLT3 or CD33 scFv joined with the EE or JUN zippers via a GS linker region, followed by a 6x His tag for purification. B. EE zipFv protein production western blot shows higher yield of FLT3 NC7 and CD33 hu195 variants compared to FLT3 D4-3 and CD33 mylo zipFvs. C. JunD zipFv protein production western blot also show higher yield of the FLT3 NC7 and CD33 hu195 variants.

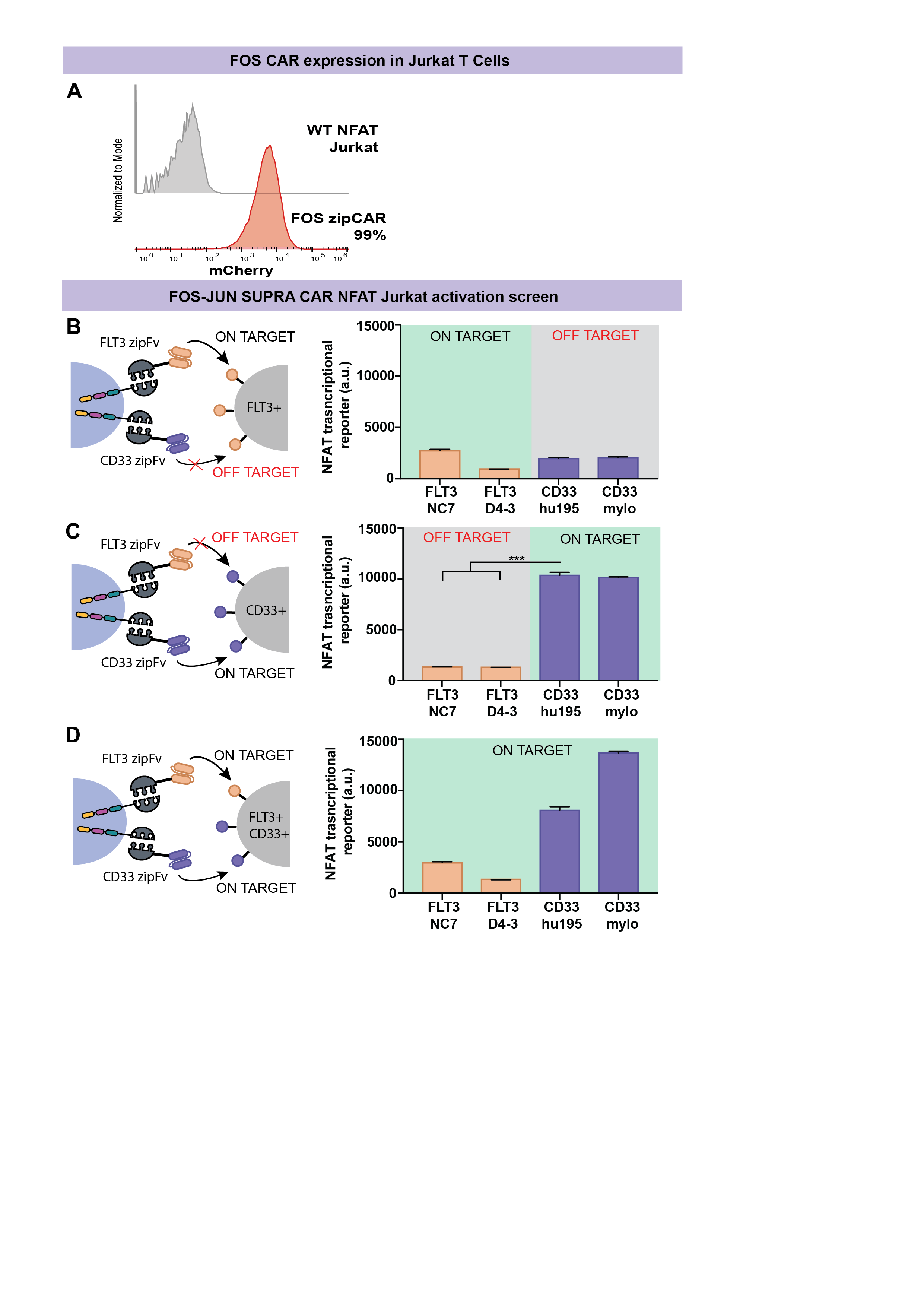

**Supplemental Figure 2. FOS-JUN SUPRA OR Gate activation screen in NFAT-GFP Jurkat cells.** A. FOS zipCAR was transduced into NFAT-GFP Jurkat cells with a transduction efficiency of 99%. B. FLT3+ K562 cells were co-cultured with FOS CAR in the presence of FLT3 JunD zipFv to assess on-target activity or CD33 JunD zipFv to assess off-target activity. There was no significant difference between on-target and off-target activity for FLT3. C. CD33+ K562 cells were co-cultured with FOS CAR in the presence of CD33 JunD zipFvs to assess on-target activity or FLT3 JunD zipFvs to assess off-target activity and showed significant on-target activation for both CD33 zipFvs. D. Double+ CD33+ FLT3+ K562 were co-cultured with FOS CAR in the presence of each JunD zipFv to assess on target activity. CD33 zipFv variations had high activity while FLT3 zipFv variations did not. In all co-culture conditions, an E:T ratio of 1:1 was used and 50 ng of the corresponding zipFv was added. CAR-T activation is reported as expression of GFP transcribed downstream of the NFAT reporter. Values shown are the means of technical triplicate samples with error bars indicating +1 standard deviation, and p-values were calculated as described in the methods.

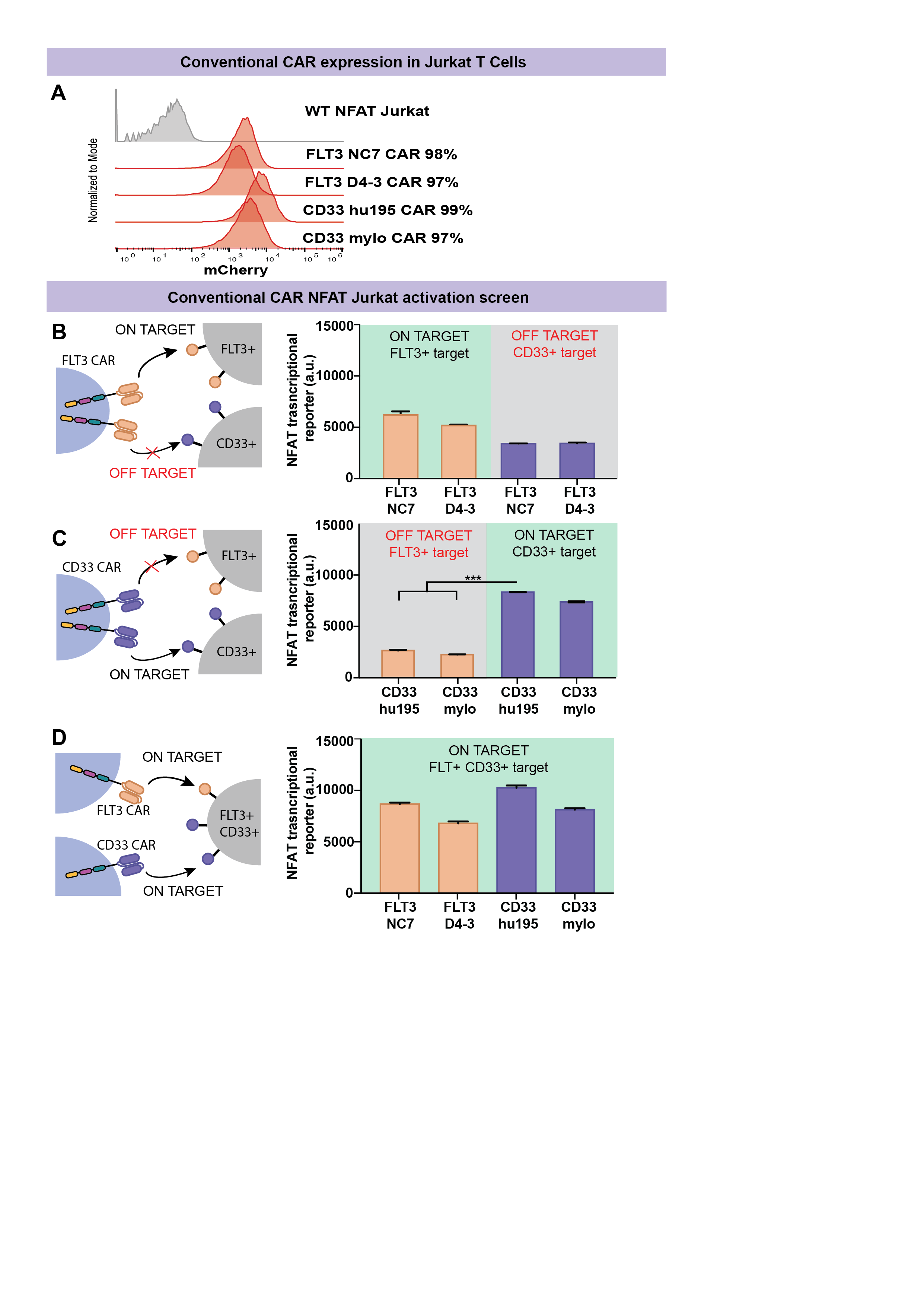

**Supplemental Figure 3. CD33 and FLT3 Conventional CAR activation in NFAT-GFP Jurkat cells.** A. Each conventional CAR was transduced into NFAT-GFP Jurkat cells with efficiencies >97%. B. FLT3 conventional CAR NFAT-GFP Jurkat cells were co-cultured with FLT3+ K562 to assess on-target activity or CD33+ K562 to assess off target activity. C. CD33 conventional CAR NFAT-GFP Jurkat cells were co-cultured with CD33+ K562 to assess on-target activity or FLT3+ K562 to assess off-target activity. D. Each conventional CAR was co-cultured with double+ CD33+ FLT3+ K562 to assess on-target activity. In all co-culture conditions an E:T ratio of 1:1 was used. CAR-T activation is reported as expression of GFP transcribed downstream of the NFAT reporter. Values shown are the means of technical triplicate samples with error bars indicating +1 standard deviation, and p-values were calculated as described in the methods.

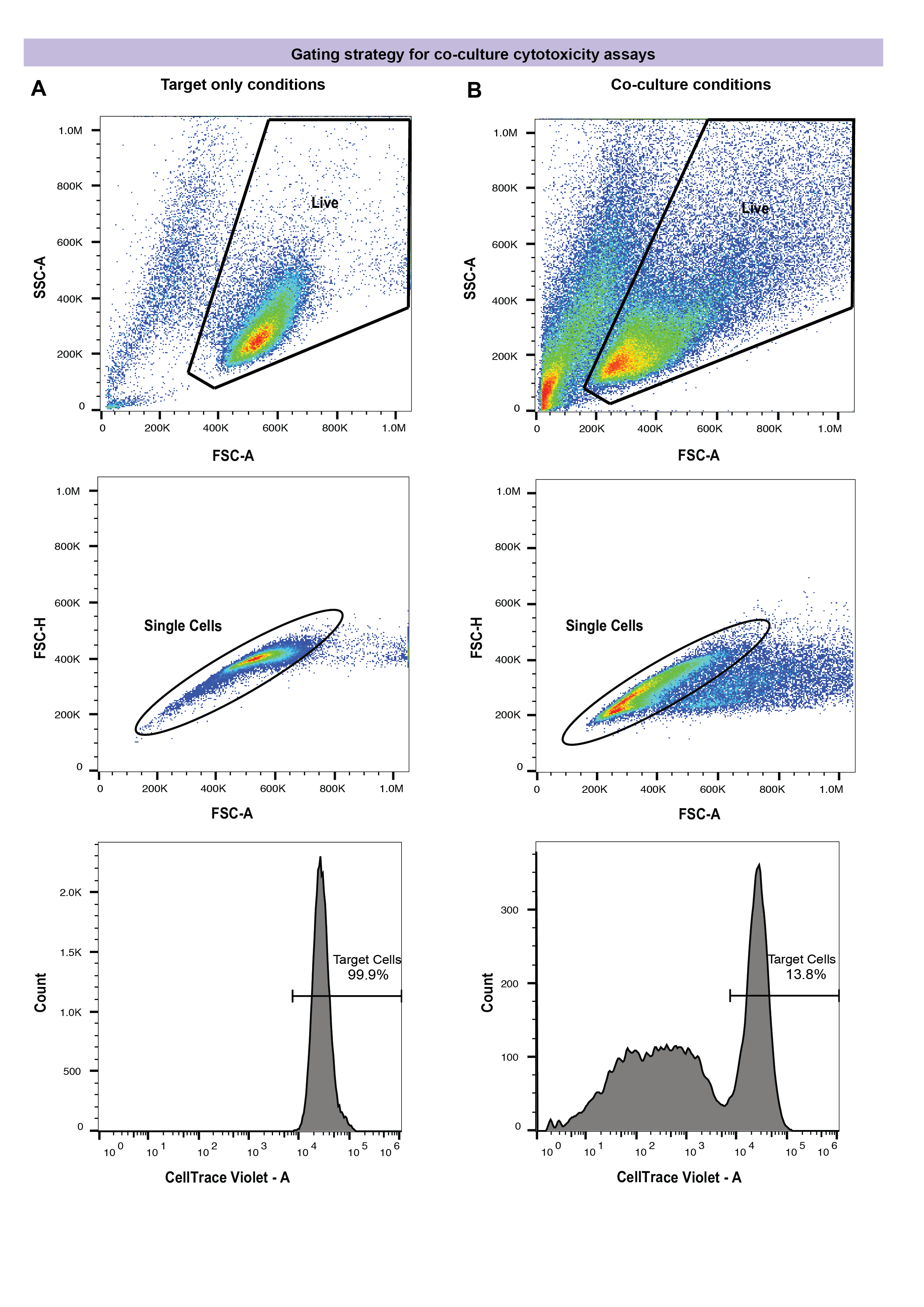

**Supplemental Figure 4. Gating strategy for co-culture cytotoxicity experiments.** A. In target only conditions, live cells were initially gated based on forward and side scatter. The live cell population was subsequently gated for single cells based on the linear region of forward scatter area versus height. Finally, target cells were shown to be positive for CellTrace Violet stain to determine the cutoff for co-culture conditions. B. In co-culture conditions, live cells were similarly gated based on forward and side scatter followed by gating of single cells based on forward scatter area versus height. Target cells were then separated from T cell populations based on CellTrace Violet to determine remaining target cell counts.

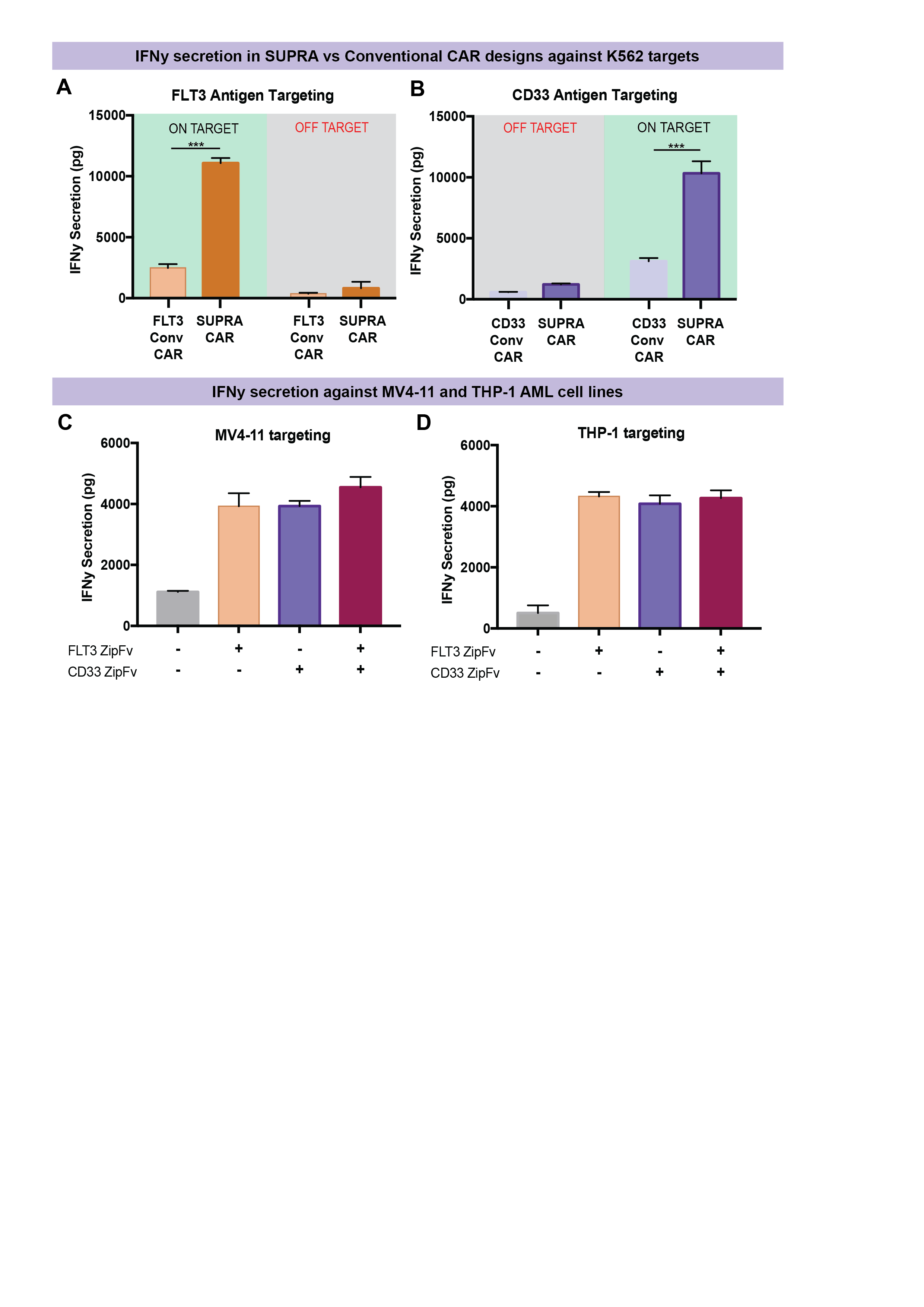

**Supplemental Figure 5. IFNy secretion from cytotoxicity assays against AML cell lines.** A. IFNy secretion by SUPRA CAR and conventional CAR transduced CD3+ T cells was assessed by ELISA and compared for on-target and off-target activation. When targeting FLT3+ cells, RR zipCAR with 100 ng FLT3 zipFv resulted in significantly higher IFNy secretion than conventional FLT3 CAR at the same E:T ratio. B. When targeting CD33+ cells, RR zipCAR with 25 ng of CD33 zipFv resulted in significantly higher IFNy secretion than CD33 conventional CAR at the same E:T ratio. C. RR zipCAR was co-cultured with MV4-11 target cells at an E:T ratio of 2:1 in the presence or absence of 125 ng of FLT3 zipFv and 25 ng CD33 zipFv. IFNy secretion was assessed via ELISA and shows high secretion of IFNy when either or both zipFvs were present compared to no zipFv conditions. D. RR zipCAR was co-cultured with THP-1 target cells at an E:T ratio of 2:1 in the presence of absence of 125 ng FLT3 zipFv and 25 ng CD33 zipFv. IFNy secretion was assessed via ELISA and shows high secretion of IFNy when either or both zipFvs were present compared to no zipFv conditions.

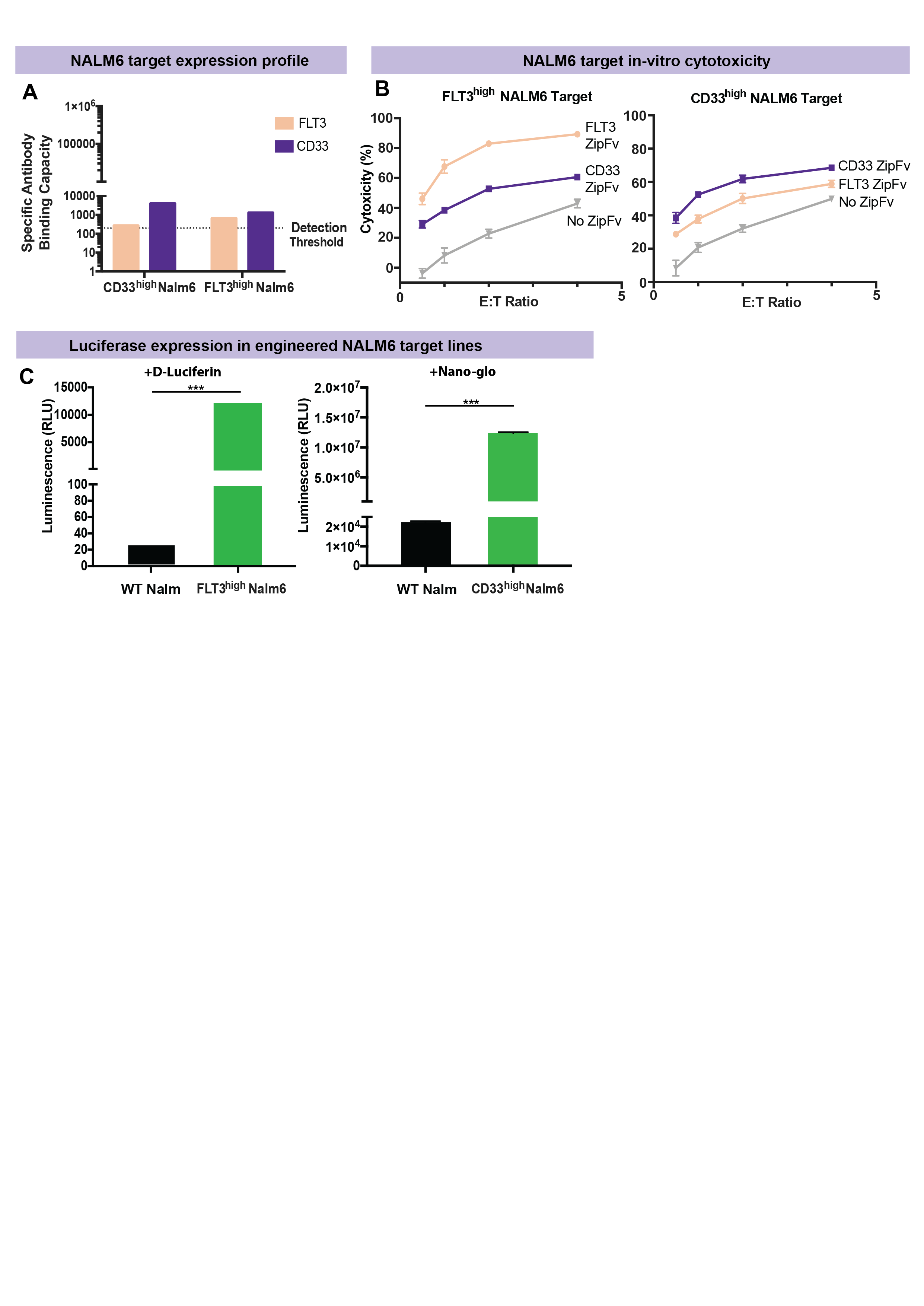

**Supplemental Figure 6. Validation of engineered NALM6 cell lines for *in vivo* studies.** A. MESF quantitative fluorescence kit from Bangs Laboratories was used to quantify the antibody binding capacity against FLT3 and CD33 antigens on engineered Nalm6 cell lines to confirm variable expression profiles. B. RR zipCAR was co-cultured *in vitro* with the engineered Nalm6 cell lines in the presence or absence of 100 ng FLT3 zipFv and 25 ng CD33 zipFv at various E:T ratios to assess specific lysis of each target cell line prior to *in vivo* injection. C. To enable imaging of each cell line separately in the same mouse, FLT3^high^ Nalm6 were engineered to express Firefly luciferase, confirmed by luminescence imaging after the addition of D-luciferin, and CD33^high^ Nalm6 were engineered to express Antares luciferase, confirmed by luminescence imaging after the addition of Nano-glo substrate.

**Supplemental Figure 7. Scaling *in vitro* zipFv dosage for preliminary *in vivo* trials**

In vitro assay conditions given an E:T ratio of 1:1

$$\left[ T cells \right]=\frac{25k}{200 uL}=\frac{125k}{mL}$$

$$\left[ Nalm6 \right]=\frac{25k}{200 uL}=\frac{125k}{mL}$$

ZipFv dose where we start to see Hook effect with CD33 zipFv is 25 ng/well

$$\left[ zipFv \right]=\frac{25 ng}{200 uL}=\frac{0.125 ug}{mL}$$

For in vivo trial experiments, 500k Nalm6 are injected on D0. Assuming a 24 hour doubling rate beginning D1, we can expect approximately 8-10 million cells by D6. We also assume ~2 mL total blood volume per mouse.

$$\left[ T cells \right]=\frac{5 mil}{2 mL}=\frac{2.5 mil}{mL}$$

$$\left[ Nalm6 \right]=\frac{10 mil}{2 mL}=\frac{5 mil}{mL}$$

$$\frac{\left[ In vivo density \right]}{\left[ In vitro density \right]}=\frac{\frac{7.5mil}{mL}}{\frac{0.25mil}{mL}}= 30X$$

We can scale zipFv dose based on this 30X increase in cell density

$$\frac{0.125ug}{mL}\times30\times2mL=7.5\frac{ug}{mouse}$$

$$\boldsymbol{Assuming \sim25g mouse, this equates to a dose of 0.3}\frac{\boldsymbol{mg}}{\boldsymbol{kg}}$$

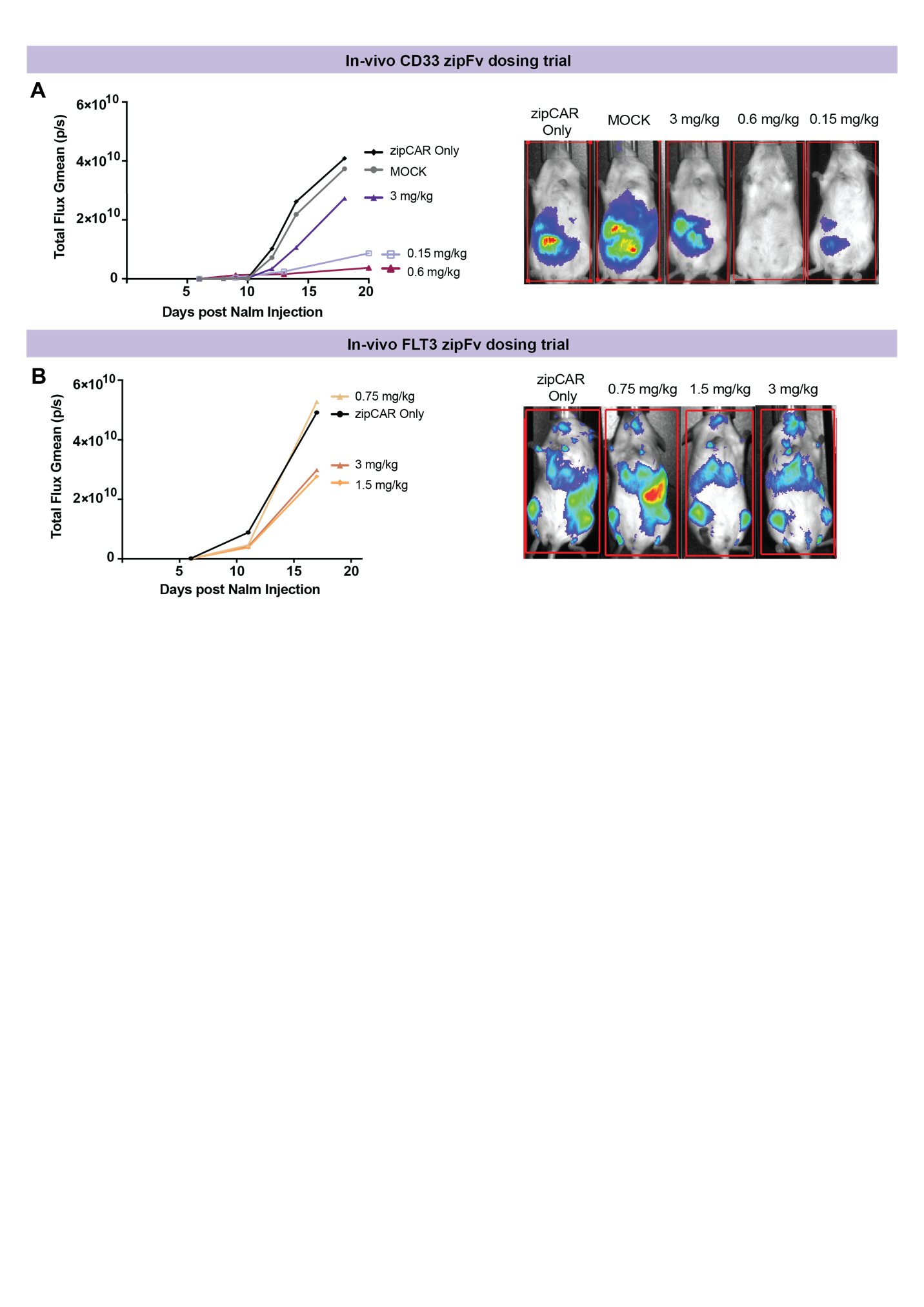
**Supplemental Figure 8. Preliminary *in vivo* dosing trials of FLT3 and CD33 zipFvs.** A. To determine a zipFv dose for CD33, two mice per group were injected with CD33^high^ Nalm6, followed by zipCAR and either no zipFv or 4 doses of 0.15 mg/kg, 0.6 mg/kg, or 3 mg/kg of CD33 zipFv. Total flux upon addition of FFz is plotted as Gmean to account for variation due to the small sample size. B. To determine the zipFv dose for FLT3, two mice per group were injected with FLT3^high^ Nalm6, followed by zipCAR and either no zipFv or 4 doses of 0.75 mg/kg, 1.5 mg/kg, or 3 mg/kg of FLT3 zipFv. Total flux upon addition of D-luciferin is plotted as Gmean to account for variation due to the small sample size.
